# Trabectedin derails transcription-coupled nucleotide excision repair to induce DNA breaks in highly transcribed genes

**DOI:** 10.1101/2023.07.10.548294

**Authors:** Kook Son, Vakil Takhaveev, Visesato Mor, Hobin Yu, Emma Dillier, Nicola Zilio, Nikolai J.L. Püllen, Dmitri Ivanov, Helle D. Ulrich, Shana J. Sturla, Orlando D. Schärer

**Affiliations:** Center for Genomic Integrity, Institute for Basic Science (IBS), 44919 Ulsan, Republic of Korea; Department of Health Sciences and Technology, ETH Zürich, 8092 Zürich, Switzerland; Department of Biological Sciences, Ulsan National Institute of Science and Technology (UNIST), 44919 Ulsan, Republic of Korea; Institute of Molecular Biology (IMB), 55128 Mainz, Germany

**Author notes:** These authors contributed equally (Kook Son, Vakil Takhaveev). Corresponding Authors: Orlando D. Schärer, Shana Sturla.

## Abstract

Most genotoxic anticancer agents fail in tumors with intact DNA repair. Therefore, trabectedin, a unique agent more toxic to cells with active DNA repair, specifically transcription-coupled nucleotide excision repair (TC-NER), provides new therapeutic opportunities. To unlock the potential of trabectedin and inform its application in precision oncology, a full mechanistic understanding of the drug’s TC-NER-dependent toxicity is needed. Here, we determined that abortive TC-NER of trabectedin-DNA adducts forms persistent single-strand breaks (SSBs) as the adducts block the second of the two sequential NER incisions. We mapped the 3’-hydroxyl groups of SSBs originating from the first NER incision at trabectedin lesions, recording TC-NER on a genome-wide scale. We showed that trabectedin-induced SSBs primarily occur in transcribed strands of active genes and peak near transcription start sites. Frequent SSBs were also found outside gene bodies, connecting TC-NER to divergent transcription from promoters. This work advances the use trabectedin for precision oncology and for studying TC-NER and transcription.

## INTRODUCTION

Numerous anticancer agents, including cisplatin, exert their therapeutic effects by inducing DNA damage and subsequently inhibiting essential cellular processes such as DNA replication and transcription. These agents do not work well in cancers with intrinsically high DNA repair or upregulated DNA repair as a response to chemotherapy ^1, 2^. Trabectedin (also called ET743), an antitumor drug used for the treatment of sarcoma and ovarian cancer ^3, 4^, is an unusual case, as it is more potent in certain DNA-repair-proficient cells ^5, 6^. Derived from the sea squirt *Ecteinascidia turbinata,* trabectedin is a complex natural product known to form adducts with DNA at the *N*^2^-position of dG (**Fig. 1a**) ^7–9^. The unique mechanism of trabectedin toxicity stems from the fact that cells with transcription-coupled nucleotide excision repair (TC-NER) defects exhibit resistance to the drug, while TC-NER-proficient cells accumulate DNA breaks following treatment ^5, 10^. This suggests that the breaks formed by TC-NER upon trabectedin treatment are more toxic than the original DNA adduct. Although earlier studies on the properties of trabectedin have provided clinically relevant insights for its use, the understanding of the exact mechanisms underlying its toxicity remains limited, restricting the drug’s application in precision medicine.

A key to the mechanism of trabectedin toxicity lies in understanding of its interaction with NER machinery. NER operates through two subpathways: global genome (GG)-NER and TC-NER ^11^. GG-NER is initiated by damage sensors XPC-RAD23B and UV-DDB (DDB1-DDB2 complex), which recognize lesions that induce thermodynamic destabilization in a DNA duplex ^12^. In contrast, TC-NER is initiated by stalling of an RNA polymerase and driven by CSB, CSA, UVSSA, and ELOF1 ^13^. Though the trabectedin-DNA adduct is bulky and causes a minor bend in the DNA ^9, 14^, it has a thermodynamically stabilizing effect on the DNA duplex ^15, 16^, which is consistent with the lack of recognition by GG-NER ^14, 17^. Therefore, trabectedin adducts can only be acted upon by the TC-NER pathway (**Fig. 1b**). A critical unsolved question is how trabectedin induces TC-NER-dependent breaks in DNA, especially given that other TC-NER-specific DNA lesions, for example those formed by illudin S and acylfulvene, undergo complete repair ^18–21^.

**Fig.1:**
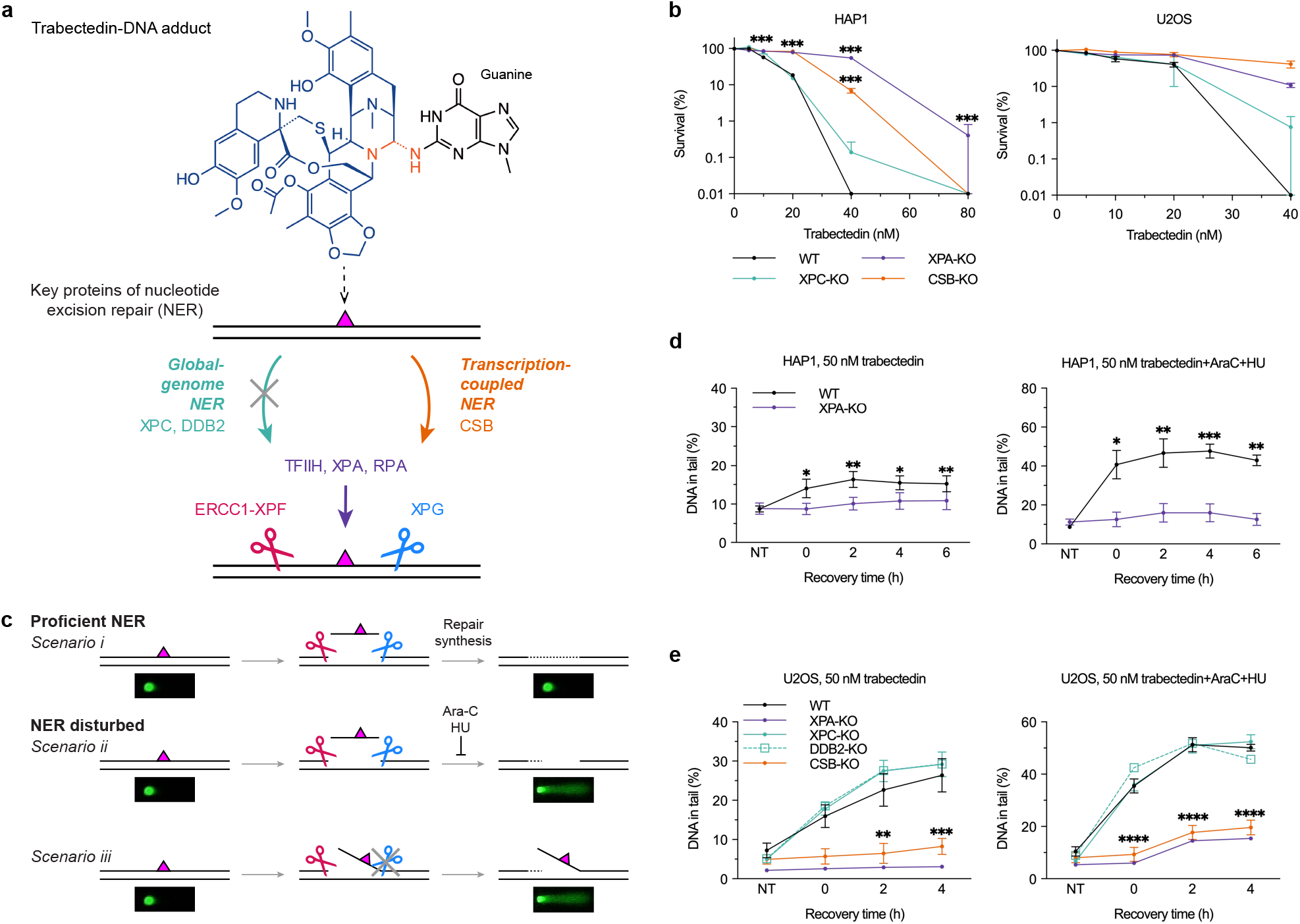
Trabectedin induces TC-NER-dependent DNA strand breaks in G1 cells. **a**, Structures of trabectedin and trabectedin-DNA adduct. These adducts are recognized by TC-NER, but not GG-NER. **b**, HAP1 or U2OS WT, XPC-, XPA-, and CSB-KO cells were treated with the indicated doses of trabectedin or DMSO control for 2 h and incubated with fresh medium. The number of colonies was counted after 8 days. Mean ± SEM of 3 biological replicates (3 technical replicates per experiment) for HAP1 and 2 biological replicates (3 technical replicates per experiment) for U2OS. ***P<0.001 using ordinary two-way ANOVA with Dunnett’s multiple comparisons test (between WT and mutants at each concentration). **c**, Simplified schematic of assessing NER incision activity following DNA damage by alkaline COMET chip assays. **d**, HAP1 WT and XPA-KO cells were arrested in G1 with palbociclib (2 μM, 24 h) and treated with trabectedin (50 nM, 2 h). The cells were kept in G1 and allowed to recover for up to 6 hrs with or without repair synthesis inhibitors (0.5 mM HU, 5 μM AraC). ssDNA breaks were analyzed by alkaline COMET chip assays. Mean ± SEM of 4 biological replicates. *P<0.05, **P<0.01, ***P<0.001 using two-tailed paired t-test (between WT and XPA-KO at each recovery time). **e**, U2OS WT, XPA-, XPC-, DDB2-, and CSB-KO cells were arrested in G1 with palbociclib (1 µM, 24 h) and treated with trabectedin (50nM, 2 h). The cells were kept in G1 and allowed to recover for up to 4 hrs with or without the repair synthesis inhibitors (1 mM HU, 10 μM AraC). ssDNA breaks were analyzed by alkaline COMET chip assays. Mean ± SEM of 4 (WT, XPC-KO), 3 (CSB-KO) biological replicates. No error bars for DDB2-KO and XPA-KO (n=1). **P<0.01, ***P<0.001, ****P<0.0001 using ordinary two-way ANOVA with Dunnett’s multiple comparisons test (between WT and XPC-or CSB-KO at each recovery time).

Our goal was to explore the mechanism by which DNA single strand breaks (SSBs) form and persist following the processing of trabectedin by TC-NER. Utilizing NER-specific alkaline COMET chip assays ^22^ and various mutant cell lines, we conducted a systematic analysis of TC-NER-dependent break induction following trabectedin treatment. In NER, damage is removed through a dual incision reaction, first by ERCC1-XPF on the 5’ side to the lesion, followed by XPG on the 3’ side to the lesion ^23^. Our findings indicate that while XPF incision occurs normally, the catalytic activity of XPG is inhibited by trabectedin-DNA adducts. We leveraged this newly discovered mechanism of trabectedin-induced SSB formation to map XPF-mediated incision sites, revealing TC-NER activity on a genome-wide scale as well as suggesting that XPF may cleave DNA in a sequence-specific way. Our analysis showed that trabectedin induces SSBs predominantly on the transcribed strand of active genes and to a lesser degree on the opposite strand upstream of gene bodies due to divergent transcription. Characterizing trabectedin-induced SSB landscapes across diverse genotypes, we developed a robust approach – which we call TRABI-seq – for probing TC-NER as well as transcription. The mechanistic insight from our research could advance trabectedin’s use in precision oncology with trabectedin serving both as a drug and a diagnostic for functional characterization.

## RESULTS

### Trabectedin induces TC-NER-dependent DNA strand breaks in G1 cells

TC-NER deficiency renders cells resistant to trabectedin (**Fig. 1a**) but sensitive to illudin S ^5, 6, 18, 20, 21^. To confirm the reported cytotoxicity profile, we treated TC-NER proficient or deficient HAP1, U2OS or XP patient fibroblast cell lines with trabectedin. WT HAP1 and U2OS cells as well as XPA-mutant patient cells (XP2OS; XP-A) complemented with XPA-WT showed an IC50 in the range of 20-30 nM in clonogenic survival assays (**Fig. 1b, Supplementary Fig. 1a**). GG-NER-deficient XPC-knockout HAP1 and U2OS cells were as sensitive as the WT cells, while cells deficient in the TC-NER factor CSB were strikingly resistant (**Fig. 1b**). Similarly, resistance was observed for XPA knockout HAP1, U2OS as well as XPA-mutant XP2OS cells; these three cell lines are defective in both NER pathways (**Fig. 1b**, **Supplementary Fig. 1a**). By contrast, and consistent with previous reports ^18, 21^, HAP1 cells deficient in CSB and XPA displayed marked sensitivity to illudin S, a regular TC-NER substrate, compared to WT and XPC-deficient cells (**Supplementary Fig. 1b**).

We hypothesized that the transcription-stalling trabectedin-DNA adducts undergo an abortive TC-NER reaction that results in the formation of persistent SSBs. To test this hypothesis, we set out to detect SSBs using high throughput alkaline COMET chip assays^22^. In a standard NER reaction, for example following UV damage induction, SSBs are transiently formed after incision of the damaged strand and before completion of repair synthesis and ligation. Under normal conditions, gaps formed by removal of lesions are very short-lived and not revealed by COMET assays (**Fig. 1c**, *scenario i*). By contrast, if NER reactions are carried out in the presence of DNA repair synthesis inhibitors AraC/HU, persistent SSBs are formed ^24, 25^ (**Fig. 1c**, *scenario ii*). We verified that these SSBs can be readily detected by COMET Chip assays in UV-exposed XP2OS cells and that comet tails were only detected if the cells expressed WT-XPA and AraC/HU were added (**Supplementary Fig. 1c**).

Having established a robust method for break detection, we tested if trabectedin induced SSBs in a TC-NER dependent manner (**Fig. 1c**, *scenario iii*). To reduce the background signal of breaks formed during replication, we synchronized the cells in G1 using the CDK4/6 inhibitor palbociclib. We first treated HAP1 cells with trabectedin, and measured time-dependent break formation by the COMET chip assay. Treatment with trabectedin alone resulted in break formation about two-fold over background and the signal was increased to about four-fold in the presence of Ara-C/HU (**Fig. 1d**, **Supplementary Fig. 1d**). No breaks were observed in XPA-deficient cells, showing that the break formation was NER-dependent (**Fig. 1d, Supplementary Fig. 1e**). In WT U2OS cells, breaks were increased about four-fold 4 h after treatment with trabectedin in the absence of DNA synthesis inhibitors (**Fig. 1e, Supplementary Fig. 1f**). Breaks were formed to a similar extent in XPC-and DDB2-deficient cells, showing that these breaks are not caused by GG-NER (**Fig. 1e, Supplementary Fig. 1g-h**). Breaks were absent in CSB-and XPA-deficient cells, in line with TC-NER being responsible for break formation (**Fig. 1e, Supplementary Fig. 1i-j**). In U2OS cells, trabectedin-induced breaks were again increased in the presence of AraC/HU (from ∼30% to ∼50% DNA in the tail, albeit with a slightly higher background signal in AraC/HU-treated cells) (**Fig. 1e, Supplementary Fig. 1f-j**). Our results demonstrate that trabectedin induces DNA breaks in a TC-NER dependent manner in G1 cells.

### XPF catalytic activity is necessary for trabectedin-induced DNA breaks

In NER, the incision 5’ to the lesion by the XPF-ERCC1 endonuclease precedes the incision 3’ to the lesion by the XPG endonuclease ^23^. Therefore, we hypothesized that the XPF-mediated incision, but not the XPG-mediated incision, is required for trabectedin-induced break formation. To test if the 5’ incision is necessary for inducing SSBs after trabectedin treatment, we used XPF-mutant-expressing XP2YO patient fibroblasts either complemented with XPF-WT or catalytically inactive XPF-D687A as well as HAP1 cells with an XPF-D687A mutation at the endogenous locus (note that there was an additional TCG(R) deletion at 701 in endogenous XPF, which might have resulted in partial XPF protein degradation, **Supplementary Fig. 2a**). We also used ERCC1 KO cells (**Supplementary Fig. 2a**), which do not express functional XPF ^26^. In the absence of XPF catalytic activity, no breaks were observed in XP2YO cell lines expressing no or mutated XPF when treated with UV and AraC/HU (**Supplementary Fig. 2b**). The catalytic activity of XPF was necessary for trabectedin’s cytotoxicity as both ERCC1-KO and XPF-D687A cells were resistant to trabectedin in HAP1 cells (**Fig. 2a**). Similarly, expressing catalytically inactive XPF-D687A in XPF-mutant XP2YO cells did not restore sensitivity to trabectedin, while the expression of WT XPF did (**Fig. 2c)**. Next, we used COMET chip assays to assess whether the observed survival pattern correlated with break formation. SSBs were observed in WT cells treated with trabectedin as expected, but no breaks were observed in ERCC1-KO or XPF-D687A cells at 2 hours post trabectedin (**Fig. 2b**). With the caveat of the additional mutation in the HAP1 XPF-D687A cells, our results indicate that the presence and nuclease activity of the ERCC1-XPF complex are needed for trabectedin-induced break formation (**Fig. 2b and Supplementary Fig. 2c**). We note that ERCC1 and XPF are critical for genomic integrity even in the absence of exogenous damage. This role of the proteins manifests itself more in ERCC1-and XPF-deficient HAP1 versus U2OS cells, considering that we observe residual sensitivity and breaks in HAP1 cells but not in U2OS cells lacking these proteins at later time points following trabectedin treatment (**Fig. 2d**).

**Fig. 2.**
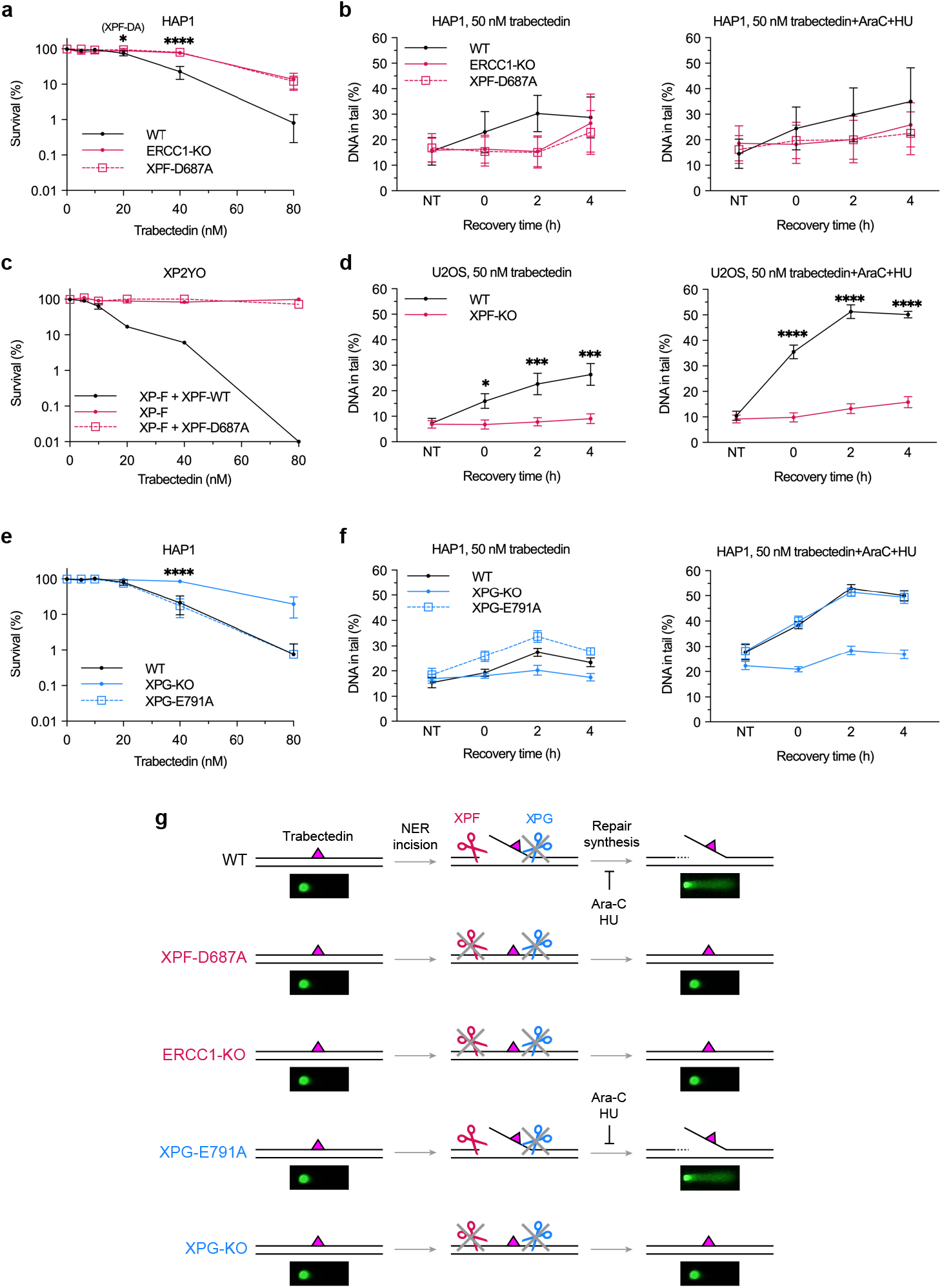
Trabectedin-induced DNA break formation and toxicity depend on the catalytic activity of XPF but not that of XPG. **a**, HAP1 WT, ERCC1-KO, and XPF-D687A cells were treated with the indicated doses of trabectedin or DMSO control for 2 h and incubated with fresh medium. The number of colonies was counted after 7 days. Mean ± SEM of 5 (WT, XPF-D687A) and 3 (ERCC1-KO) biological replicates (3 technical replicates per experiment). *P<0.05, ****P<0.0001 using ordinary two-way ANOVA with Dunnett’s multiple comparisons test (between WT and ERCC1-KO or XPF-D687A at each concentration). **b**, HAP1 WT, ERCC1-KO, and XPF-D687A cells were arrested in G1 with palbociclib (2 μM, 24 h and treated with trabectedin (50 nM, 2 h). Cells were kept in G1 and incubated for up to 4 h with or without repair synthesis inhibitors (0.5 mM HU, 5 μM AraC). ssDNA breaks were analyzed by alkaline COMET chip assays. Mean ± SEM of 5 (WT) and 3 (ERCC1-KO and XPF-D687A) biological replicates (Statistical analysis shown in Supplementary Fig. 2c). **c**, XP2YO patient cells were treated with the indicated doses of trabectedin or DMSO control for 2 h and incubated with fresh media medium. The number of colonies was counted after 8 days. Mean ± SEM of 2 biological replicates (3 technical replicates per experiment). **d**, U2OS WT and XPF-KO cells were arrested in G1 with palbociclib (1 µM, 24 h) and treated with trabectedin (50 nM, 2 h). The cells were kept in G1 and allowed to recover for up to 4 h with or without the repair synthesis inhibitors (1 mM HU, 10 μM AraC). ssDNA breaks were analyzed by alkaline COMET chip assays. Mean ± SEM of 4 biological replicates. *P<0.05, ***P<0.001, ****P<0.0001 using ordinary two-way ANOVA with uncorrected Fisher’s LSD (between WT and XPF-KO at each recovery time). **e**, HAP1 WT, XPG-KO, and XPG-E791A cells were treated with the indicated doses of trabectedin or DMSO control for 2 h and incubated with fresh medium. The number of colonies was counted after 7 days. Mean ± SEM of 4 biological replicates (3 technical replicates per experiment). ****P<0.0001 using ordinary two-way ANOVA with Dunnett’s multiple comparisons test (between WT and XPG-KO or XPG-E791A at each concentration). **f**, HAP1 WT, XPG-KO, XPG-E791A, and XPA-KO cells were arrested in G1 with palbociclib (2 µM, 24 h) and treated with trabectedin (50 nM, 2 h). Cells were kept in G1 with or without repair synthesis inhibitors (0.5 mM HU, 5 μM AraC) and incubated for up to 4 h. ssDNA breaks were analyzed by alkaline COMET chip assays. Mean ± SEM of 5 biological replicates (Statistical analysis shown in Supplementary Fig. 3d-e). **g**, Simplified schematic of assessing NER incision activity of trabectedin DNA adducts in HAP1 WT, XPG-E791A, and XPG-KO cells by alkaline COMET chip assays.

### Evidence that trabectedin inhibits XPG catalytic activity

Since the second incision in NER is executed by XPG endonuclease, we tested whether the catalytic activity of XPG was needed for break formation by trabectedin. For this purpose, we generated HAP1 cells with catalytically inactive XPG (XPG-E791A), (clones #22 and #45, **Supplementary Fig. 3a**), and confirmed that DNA breaks were induced following UV irradiation in the presence or absence of AraC/HU (**Supplementary Fig. 3b)**, in line with reported effects of XPG with this mutation expressed in patient cell lines ^22^. The addition of AraC/HU was not absolutely needed for the formation of SSBs following UV irradiation in cells expressing XPG-E791A, demonstrating that SSBs persist in the presence of catalytically inactive XPG (**Supplementary Fig. 3b**).

HAP1 WT or XP3BR (XP-G) patient cells complemented with XPG-WT showed the expected level of sensitivity to trabectedin, while XPG knockout HAP1 or XP3BR (XP-G) patient cells were resistant (**Fig. 2e, Supplementary Fig. 3c**). Intriguingly, XPG-E791A expressing HAP1 (clone #22) or XP3BR cells were equally sensitive to trabectedin as WT cells, showing that the catalytic activity of XPG is not needed for trabectedin-induced toxicity. The difference in sensitivity of the XPG-KO and XPG-E791A cells can be explained with the observation that the presence, but not catalytic activity of XPG, is needed for the XPF incision to occur ^23, 27^.

To determine if DNA breaks are responsible for toxicity observed in XPG-E791A cells, we conducted the alkaline COMET chip assay with XPG-WT, -KO, and -E791A cells.

There were no SSBs in XPG-KO cells at 2 h post trabectedin, whereas breaks were induced in XPG-E791A cells, seemingly at even higher levels than in WT cells in the absence of DNA synthesis inhibitors (**Fig. 2f**, left; **Supplementary Fig. 3d-e**). In the presence of DNA synthesis inhibitors, break formation was increased, and found to be at the same levels in the XPG-WT and XPG-E791A cells (**Fig. 2f**, right). The fact that trabectedin-induced break formation in WT and E791A cells is very similar supports the model that trabectedin-DNA adducts block XPG incision.

Our studies suggest that trabectedin-induced break formation and toxicity depend on the catalytic activity of XPF, but not that of XPG (**Fig. 2g**). In this way, only one NER incision is made, leading to a persistent SSBs with a free 3’-OH upstream of the trabectedin-DNA adduct.

### Trabectedin-induced DNA breaks can be mapped genome-wide

The COMET chip experiments have revealed a mechanism responsible for TC-NER-and trabectedin-induced SSB formation but provided no information on where in the genome these breaks occur. Are trabectedin-induced SSBs evenly distributed throughout the genome, do they occur in specific genomic locations, or do they perhaps target oncogenes upregulated in certain tumors?

We hypothesized that our mechanistic insight would provide a basis for an approach to reveal trabectedin-induced SSBs with single-nucleotide resolution and in a genome-wide fashion. Specifically, we aimed to map the persistent ERCC1-XPF-dependent SSBs upstream of the drug-DNA adduct, employing the recently developed method of genome-wide ligation of severed 3’-OH ends followed by sequencing, GLOE-Seq ^28^. We introduced an upgrade to the method that provides a more balanced representation of SSB signals and enables filtering out PCR amplification duplicates of sequencing reads. We labeled DNA fragments originating from DNA breaks first with a biotinylated proximal adapter for enrichment and subsequently with a distal adapter containing a unique molecular identifier (UMI) of randomized nucleotides to reflect the abundance of a given DNA break at a certain genomic position across the population of cells (**Fig. 3a**). As a positive control for the upgraded method, we treated U2OS WT DNA with the Nb.BsrDI nicking endonuclease that introduces SSBs before CATTGC sequences and confirmed that GLOE-Seq maps breaks at this sequence with at least 82-84% precision, *i.e.*, the fraction of reads revealing the CATTGC pattern, (**Supplementary Fig. 4a**) and 87-88% sensitivity, *i.e.*, the fraction of identified Nb.BsrDI sites in the human reference genome (**Supplementary Fig. 4b**). This analysis showed that the upgraded method performed on the human genome at least as well as the original GLOE-Seq protocol, which was validated in the same way in the yeast genome ^28^.

**Fig. 3.**
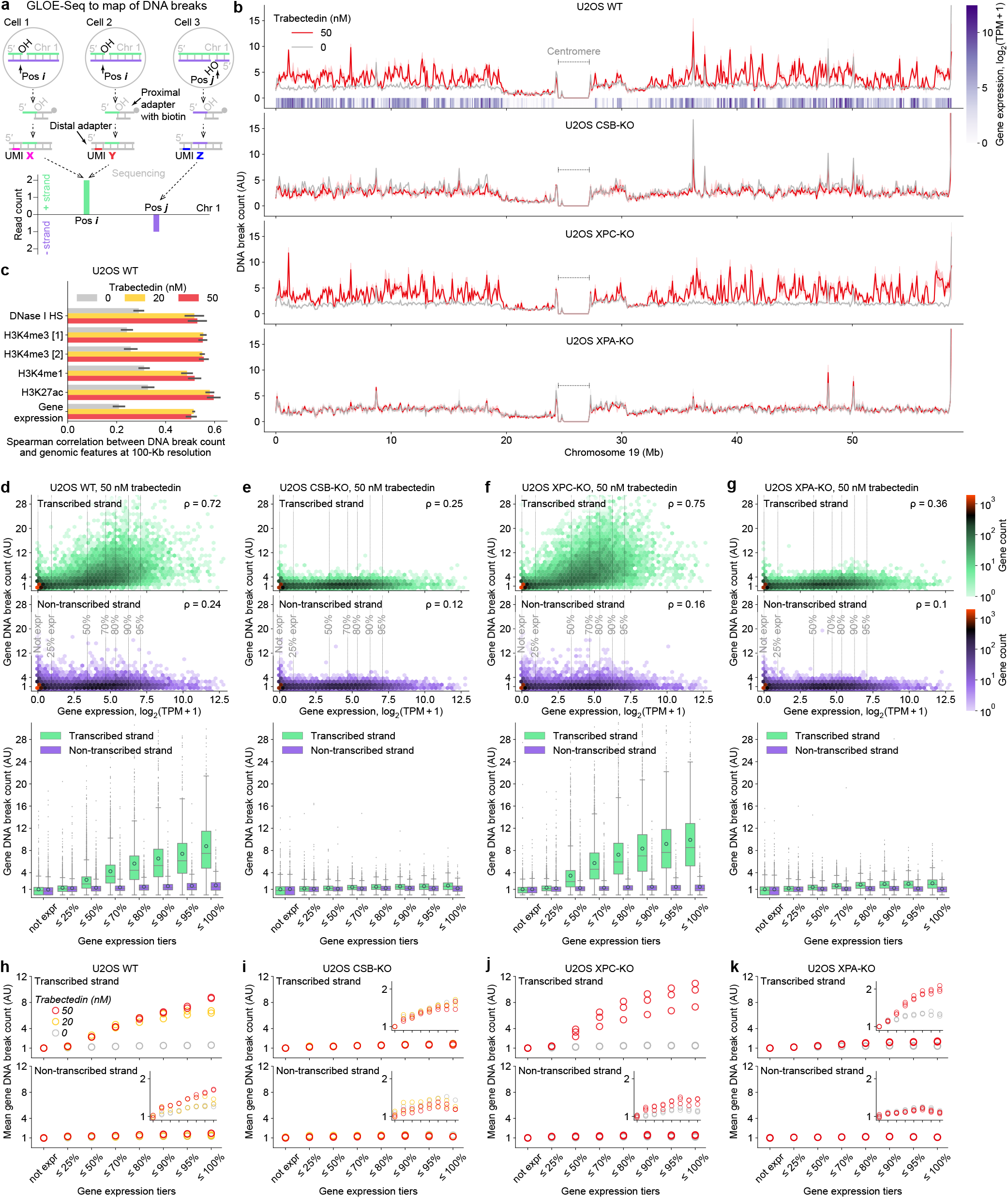
Trabectedin-induced DNA break counts correlate with gene expression levels. **(a)** Principle of GLOE-Seq that maps DNA breaks in a genome-wide and strand-specific fashion and provides an estimate of the frequency of individual breaks in a population of cells. The distal adapter with a unique molecular identifier (UMI) is an upgrade from the original GLOE-Seq protocol ^28^. Pos: position, chr: chromosome, OH: free 3’ hydroxyl. **(b)** DNA break count along chromosome 19 in four cell lines after two-hour exposure to trabectedin or vehicle (DMSO) and subsequent 2 h recovery. Solid line and shaded area: mean ± s.d. across biological replicates. Vertical bar heatmap: gene expression level in unexposed U2OS WT. All data are shown per 100-Kb bin. **(c)** Genome-wide correlation (shown as mean ± s.d. across replicate experiments) of DNA break count with the abundance of DNase I hypersensitivity (HS) sites (transcriptional activity), H3K4me3 (active gene promoters), H3K4me1 (active enhancers) and H3K27ac (active promoters and enhancer) as well as gene expression. **(d-g)** DNA break count on each strand of protein-coding genes in U2OS WT (d), CSB-KO (e), XPC-KO (f) and XPA-KO (g) after 2 h exposure to trabectedin and subsequent 2 h recovery versus gene expression level in unexposed U2OS WT. ρ: Spearman correlation coefficient calculated for continuous (not tiered) data of gene expression and DNA break count (correlation analysis for all replicates in **Supplementary Fig. 5h**). Heatmaps: the color of a hexagonal bin reflects the gene count. Box plots: gene expression was categorized in tiers via percentiles shown by vertical lines in the heatmaps; boxes are interquartile ranges, internal horizontal lines are medians, circular markers are means, whiskers extend to the furthest datapoint within 1.5x interquartile range, datapoints beyond are shown as small markers. Number of genes per tier and number of genes beyond the maximal y-axis value are provided in **Methods**. Presented data: one biological replicate of drug exposure per cell line. **(h-k)** Mean DNA break count of protein-coding genes in U2OS WT (h), CSB-KO (i), XPC-KO (j) and XPA-KO (k) after 2 h exposure to trabectedin or vehicle and subsequent two-hour recovery versus gene expression level in unexposed U2OS WT. Presented data: multiple biological replicates. Gene expression tiers are described in (d-g). The internal plots zoom in on the y-axis. b, d-g: TPM, transcripts per million transcripts. b, d-k: AU, **Methods** describe DNA break count normalization.

Using the upgraded GLOE-Seq method, we set out to reveal the genomic positions of the ERCC1-XPF-mediated trabectedin-induced breaks. We determined genome-wide profiles of DNA breaks in the TC-NER-proficient WT U2OS cell line, the corresponding TC-NER-deficient CSB knockout, global-genomic-NER-deficient XPC knockout and fully NER-deficient XPA knockout cells. Cells were treated with trabectedin (50 nM) or vehicle control using the same conditions as in the COMET-chip assays (two-hour treatment and two-hour recovery). We found abundant trabectedin-induced DNA breaks in the TC-NER-proficient cells (U2OS WT and XPC-KO), while there were no trabectedin-induced DNA breaks in the TC-NER-deficient cells (CSB-KO and XPA-KO), as shown in detail for Chromosome 19 (**Fig. 3b**) and in an overview for the whole genome (**Supplementary Fig. 4c**). These GLOE-Seq results were consistent across biological replicates (Spearman correlation coefficient mostly around 0.9) (**Supplementary Fig. 4d**). We observed that the genome-wide profile of DNA breaks in the drug-exposed U2OS WT had a marked correlation with transcriptionally active regions of chromatin (DNase I hypersensitivity sites), epigenetic marks of active promoters and enhancers (H3K4me3, H3K4me1, H3K27ac) as well as gene expression levels (**Fig. 3c**).

### Trabectedin-induced DNA break counts correlate with gene expression levels

Therefore, we next focused our analysis on genes, and plotted the DNA break counts (Y axis) for transcribed (green) and non-transcribed strands (purple) versus gene expression levels (X axis) (**Fig. 3d-g**). We normalized the DNA break count data in each experiment by fixing the average DNA break count for transcribed (antisense) strand of unexpressed genes to 1. Gene DNA counts were normalized by gene length as longer genes naturally have more breaks (see normalization formulas in ***Methods***). We discovered that in U2OS WT cells, for the vast majority of unexpressed genes and genes up to the 50th percentile of gene expression, the DNA break count in the transcribed strand was below 4. By contrast, a more pronounced increase of DNA break count was observed at higher gene expression levels (**Fig. 3d**, heatmap). When we stratified the genes based on expression levels, we found that the average break count in the transcribed strands of the top 5% expressed genes was around 9-fold higher than in unexpressed genes, whereas DNA break count in the non-transcribed (sense) strand did not similarly depend on gene expression (**Fig. 3d**, boxplot). These data show that trabectedin induces DNA breaks preferentially in transcribed strands of highly expressed genes. We observed a similar correlation in TC-NER-proficient XPC-KO cells (**Fig. 3f**) but not in TC-NER-deficient CSB-KO (**Fig. 3e**) or XPA-KO cells (**Fig. 3g**), in agreement with COMET chip assays (**Fig. 1e**). In DMSO-treated control, the break counts were the same throughout gene expression levels for all four U2OS cell lines used (**Supplementary Fig. 5a-d**). Similar patterns were observed in HAP1 WT cells (**Supplementary Fig. 5e-f**), although the fold change in break formation throughout the gene expression tiers was lower than in U2OS cells, which is consistent with the lower fold increase in the DNA break formation in HAP1 versus U2OS cells determined in the COMET-chip assay (**Fig. 1d-e**).

We found our assay to be robust, detecting a near-identical break count pattern over biological replicates (**Fig. 3h-k**, **Supplementary Fig. 5g-h**) and a lower break count at a lower trabectedin concentration of 20 nM versus 50 nM (**Fig. 3h**). Interestingly, we identified a very minor, but reproducible expression-dependent trabectedin-induced break formation on the non-transcribed strand only in TC-NER-proficient U2OS WT and XPC-KO cells (**Fig. 3h-k**, inner plots) as well as HAP1 WT (**Supplementary Fig. 5g**, inner plot). We could also detect that in the full-NER-deficient XPA-KO cells, there was still a low level (up to 2-fold) of gene-expression-dependent trabectedin-induced break formation (**Fig. 3k**, inner plots), while in the TC-NER-deficient CSB knockout, no trabectedin-induced breaks were formed (**Fig. 3i**). Besides, we observed a modest correlation between break counts and gene expression without trabectedin exposure, with this correlation for the transcribed strand being consistently higher than for non-transcribed strand (**Supplementary Fig. 5h**, gray bars), which may indicate endogenous break formation induced by transcription. While we do not currently have a biological explanation for trabectedin-induced break formation in the non-transcribed strands and in the XPA-KO cells, these observations demonstrate the quantitative nature of our assay.

### Divergent transcription is detected by trabectedin-induced DNA breaks

Having observed that trabectedin induces damage preferentially in transcriptionally active genes, we further zoomed in on the DNA break distribution throughout gene bodies and adjacent regions in the top 30% expressed genes. In U2OS WT cells treated with trabectedin (50 nM), DNA breaks were most abundant in the transcribed strand right downstream of the transcription start site (TSS), and their frequency gradually decreased throughout the gene body, being 3 to 4-fold lower at the transcription end site (TES) (**Fig. 4a**, green trace). In line with the data of **Fig. 3d-g**, this DNA break pattern was absent in TC-NER-deficient U2OS CSB-KO (**Fig. 4b**) and XPA-KO cells (**Fig. 4d**) treated with trabectedin as well as in all untreated cells (**Supplementary Fig. 6a-d,f**). It was however very similar in trabectedin-treated TC-NER-proficient U2OS XPC-KO (**Fig. 4c**) and HAP1 WT cells (**Supplementary Fig. 6e**). The peak values of DNA breaks were found around 1-2 kilobases after the TSS independently of the gene length (**Fig. 4e,f**, left panels). We surmise that the higher break formation closely downstream of the TSS is due to the stalling of RNA polymerase at the first trabectedin-DNA adducts early in the gene and the consecutive inhibition of the transcription and TC-NER activity. Elevated break levels were also detected after TES, potentially reflecting alternative longer transcripts (**Fig. 4a,c, Supplementary Fig. 6e**).

**Fig. 4.**
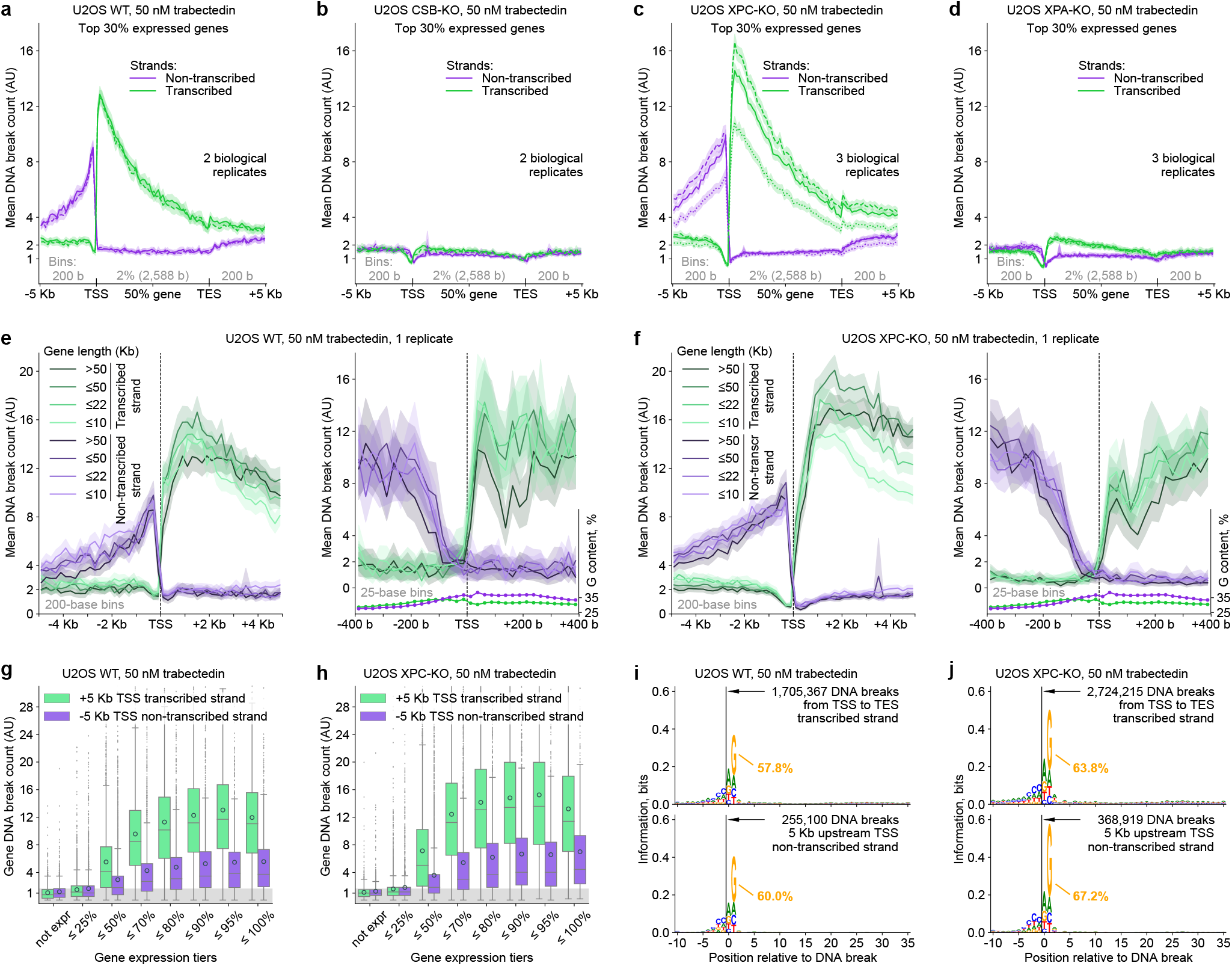
TRABI-seq detects divergent transcription and provides evidence for XPF sequence preference. **(a-d)** Strand-specific profile of the mean DNA break count throughout the gene body and its upstream and downstream 5 kilobase regions in U2OS WT **(a)**, CSB-KO **(b)**, XPC-KO **(c)** and XPA-KO **(d)** after two-hour exposure to trabectedin and subsequent two-hour recovery. 4,425 protein-coding genes, specifically, top 30% expressed in unexposed U2OS WT, are considered. Solid, dashed, and dotted curves: different biological replicates. **(e-f)** Strand-and gene-length-specific profile of the mean DNA break count in the ±5 Kb proximity of TSS in TC-NER proficient cell lines U2OS WT **(e)** and XPC-KO **(f)**, while zooming out (left panel) and in (right). The four groups of gene lengths have approximately the same number of genes. One biological replicate is shown per cell line. The same gene set as in (a-d) is considered. The strand-specific G content is calculated for the whole gene set (regardless of the gene length) using the reference genome and shows the absence of drastic sequence composition changes that would explain low DNA break counts in promoters. **(g-h)** DNA break count following two branches of divergent transcription in U2OS WT **(g)** and XPC-KO **(h)** versus gene expression. +5 Kb: within 5 Kb downstream of; -5 Kb: within 5 Kb upstream of. The plots are built analogously to Fig. 3d,f (lower panels), however, instead of the whole gene regions, the indicated 5-Kb regions are considered, which explains higher values for the transcribed strand as compared to Fig 3d,f due to non-uniform break distribution throughout the gene body (a,c). Data: one biological replicate. **Supplementary Fig. 6g-h** presents respective correlation analysis for all replicates. Gray band: threshold of endogenous DNA breaks not caused by trabectedin treatment (upper quartile of DNA break count in unexpressed genes); this threshold shows that around 25% (lower boundary of boxes) of highly expressed genes may not have trabectedin-induced breaks upstream of the TSS. **(i-j)** Sequence logos showing sequence enrichment around DNA breaks in U2OS WT **(i)** and XPC-KO **(j)**. We considered DNA breaks located in the indicated regions of the gene set used in (a-f). The percentage of G at position 1 (+2 relative to the break) is shown. Data: all biological replicates united per cell line. **Supplementary Fig. 6i-j** shows analogous analysis for TC-NER-deficient cell lines. a-f: Curve and shaded area: mean and its 95% c.i.; bin sizes are absolute (shown as a base number) or relative (shown as a percentage of gene length; the corresponding average base number indicated in parentheses). a-h: AU, **Methods** describe DNA break count normalization. a-j: TSS and TES: transcription start and end sites; Kb: kilobase; b: base.

Interestingly, we found another peak of DNA break counts a few hundred nucleotides upstream of the TSS on the non-transcribed strand, with the break count gradually decreasing yet not reaching background levels even 5 kb upstream of the TSS (**Fig. 4a,c,e,f** and **Supplementary Fig. 6e**). The frequency of the DNA breaks in the non-transcribed strand increased only further than 100 nucleotides upstream of the TSS, making this 100-nucleotide region of the promoter devoid of trabectedin-induced breaks on both DNA strands (**Fig. 4e,f**, right panels). These break patterns appear to be in line with the phenomenon of divergent transcription, *i.e.*, transcription on both DNA strands occurring in opposite directions, which is observed in most mammalian promoters ^29–32^, where the absence of strong TATA elements may lead to transcription bidirectionality ^33^. We further found that the trabectedin-induced break count upstream of the TSS in the non-transcribed strand is increasing with the gene expression level (**Fig. 4g-h**, purple; **Supplementary Fig. 6g-h**), similarly to the break count downstream of the TSS in the transcribed strand (**Fig. 4g-h**, green). However, the TTS-upstream breaks are on average two-fold less abundant than the TSS-downstream breaks in genes after the 25^th^ percentile of gene expression (**Fig. 4g-h**), which indicates that divergent transcription activity is about a half of that in the gene body averaged over the entire genome. Overall, trabectedin-induced break sequencing – a method we call TRABI-seq – reports on various types of transcription, including divergent transcription, and demonstrates that TC-NER can also occur in the intergenic space.

### TRABI-seq may reveal a sequence preference for XPF incision

As the COMET assay experiments showed that trabectedin-induced DNA breaks are caused by ERCC1-XPF activity (**Fig. 2b**), locating these breaks in TRABI-seq provides an opportunity to explore potential sequence preferences of the incision activity of this endonuclease. To analyze whether the ERCC1-XPF has a sequence preference, we focused on the top 30% expressed genes and examined DNA breaks in the regions most affected by the drug, namely, the transcribed strands of the genes and 5 kilobases of the non-transcribed strands upstream of the TSS, where any sequence preference would not be obscured by background signals (**Fig. 4a,c**). We aligned the sequence contexts of these breaks and discovered that there is an enrichment of guanine at the second position downstream of DNA breaks (**Fig. 4i-j**). According to these data, ERCC1-XPF may prefer a guanine (or complementary cytosine) one base downstream of the incision site. No overrepresented nucleotides were found further downstream of the break location (**Fig. 4i-j**), or in TC-NER deficient CSB-KO or XPA-KO cells (**Supplementary Fig. 6-j**). Thus, by providing evidence for ERCC1-XPF DNA sequence specificity, TRABI-seq serves as a tool to study the fundamental properties of NER.

## Discussion

DNA damaging agents are crucial therapeutics used in the treatment of most cancers, however, innate or acquired resistance of tumors to these agents is a major limitation. New developments to increase the efficacy of anticancer therapy rely on precision oncology where specific drugs are matched with the genetic profile of tumors. As successful clinical examples, cisplatin and poly (ADP-ribose) polymerase inhibitors (PARPi) target tumors with defects in BRCA1/2 and other genes involved in mediating homologous recombination and counteracting replication stress. Similarly, tumors with defects in NER are also hypersensitive to cisplatin, while high NER activity is associated with resistance. Recent studies have additionally put forward irofulven, a derivative of illudin S, as an agent to selectively target tumors with deficiencies in TC-NER ^34, 35^. The therapeutic action of PARP inhibitors, cisplatin and irofulven thus relies on synthetic lethality, *i.e.*, tumor vulnerability due to a defect in the repair pathway counteracting the toxic effects of the drug.

Here we investigated trabectedin, a drug that shows an opposite mode of action, namely being more toxic to cells with high repair capacity. Trabectedin is a promising drug to treat tumors with high TC-NER activity or, generally, with intact DNA repair machinery. Additionally, trabectedin may be valuable to overcome the NER-mediated resistance of tumors to cisplatin. To further its use in precision medicine, we set out to study its mechanism of NER-induced toxicity. Using highly sensitive COMET chip assays, we show that trabectedin induces persistent DNA breaks in a TC-NER dependent manner in cells that are hypersensitive to the drug. Our data suggests a model that the trabectedin-DNA adducts block the incision of the XPG endonuclease, which results in persistent XPF-mediated breaks (**Fig. 5**). We mapped those breaks genome-wide and showed that they form predominantly in highly transcribed genes and their upstream regions in association with divergent transcription. This approach of break sequencing, which we call TRABI-Seq, is informed by the uncovered mechanism of trabectedin and provides opportunities to study TC-NER and profile tumor vulnerability by mapping TC-NER activity.

**Fig. 5.**
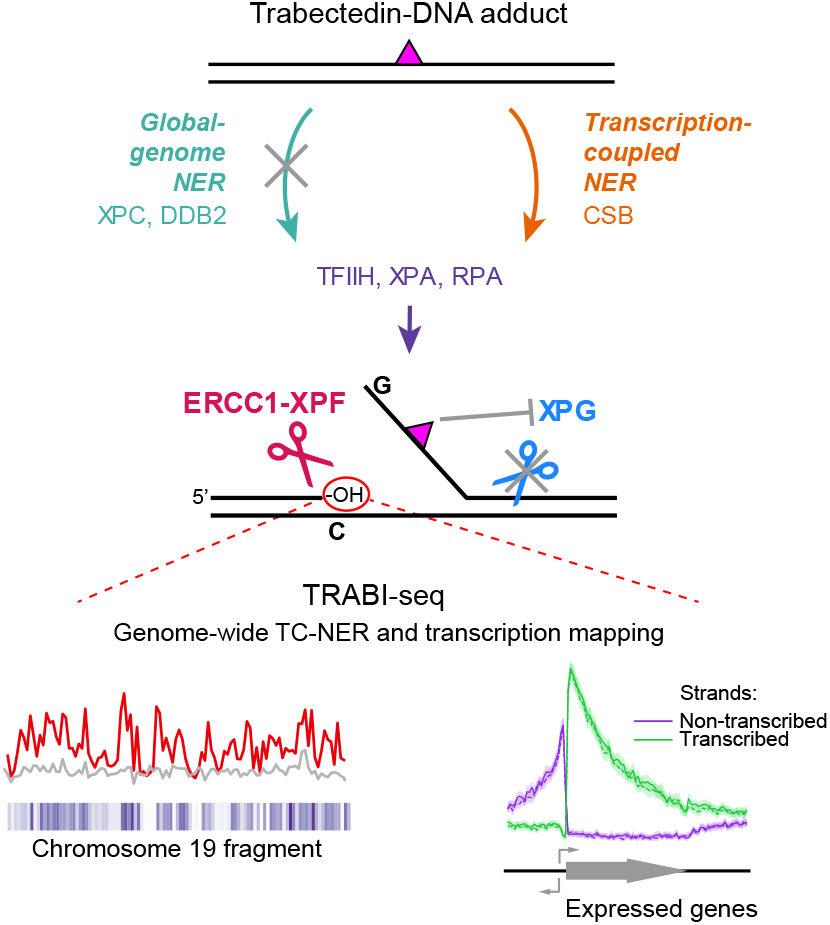
Summary of the mechanism of trabectedin-induced TC-NER-mediated SSB formation and toxicity, and the development of TRABI-Seq.

The discovered mechanism of trabectedin action is in line with the prevailing model for NER dual incision where XPF cuts before XPG ^23, 36^. Our data shows that during TC-NER of trabectedin, 5’ incision by XPF occurs normally, while 3’ incision by XPG is blocked. This observation is consistent with the fact that the incisions in NER are asymmetric with respect to the position of lesions being located much closer to the 3’ incision site by XPG (2-8 bases) than to the 5’ incision site by ERCC1-XPF (15-24 bases) ^37–39^. Furthermore, structural models indicate that the bulky trabectedin DNA adduct is oriented toward the 3’ incision site, where it may interfere with the catalytic activity of XPG^9^.

A potential application of trabectedin in precision medicine could be profiling tumors for TC-NER activity via TRABI-seq. Validated in a range of knock-out cell lines (**Fig. 3h-k**, **Supplementary Fig. 5g-h**), this robust and sensitive assay could functionally identify TC-NER deficiency in tumors, which, for instance, would be an indication for the drug irofulven, which is synthetic lethal in combination with inactive TC-NER ^34, 35^. TRABI-seq-based reporting of high TC-NER activity in tumors would, on the contrary, suggest trabectedin as a therapeutic. Along with profiling TC-NER activity, TRABI-seq could also characterize tumors with respect to their oncogene expression patterns (**Supplementary Fig. 7**) as gene break counts strongly correlate with gene expression (**Fig. 3d,h**). Interestingly, a recent report used a biotinylated derivative of lurbinectedin, a close relative of trabectedin, to map lurbinectedin-DNA adducts in a Chip-seq approach ^40^. While this method can locate the adducts to the promotors of tumor-driving genes, TRABI-seq reports also on the location of cytotoxic breaks, TC-NER and gene expression.

The persistent DNA breaks resulting from trabectedin-adduct-induced recruitment and consecutive abortion of TC-NER machinery offer a unique opportunity to study this pathway of DNA repair. By using TRABI-Seq, we surprisingly discovered that TC-NER is active beyond genes, likely due to divergent transcription branch oriented in the direction upstream of promoters and using the non-transcribed (from the gene perspective) DNA strand as a template (**Fig. 4a,c**, **Supplementary Fig. 6e**) ^29–32^. Another intriguing aspect of TRABI-Seq data is evidence for DNA sequence guiding XPF activity (**Fig. 4i-j**), which requires further investigation but is in line with recent findings regarding another DNA-repair endonuclease, MRE11-RAD50-XRS2, which was also found to cleave with a sequence preference ^41^.

In conclusion, we uncovered that trabectedin blocks one of two incision reactions in TC-NER and used this finding to map TC-NER and gene expression activity on genome-wide scale. This insight and developed techniques will advance investigations into the mechanism of trabectedin toxicity and inform its use in precision oncology.

## METHODS

### Trabectedin, illudin S and antibodies

Trabectedin used in survival and COMET experiments was from Tecoland and in GLOE-seq experiments from Lucerna-Chem AG. Illudin S was from MGI Pharma. β-actin (catalog no. MA5-15739) was from Invitrogen. XPG (catalog no. A301-484A) from Bethyl and XPF (catalog no. ab76948) from Abcam. ERCC1 (D-10) (catalog no. Sc-17809) was from Santa Cruz Biotechnology.

### Cell lines

CML derived human HAP1 wild-type, XPC-, XPA-, CSB-and XPG-KO cells were from Horizon Discovery. HAP1 XPG-E791A, XPF-D687A, and ERCC1-KO cells were generated with CRISPR-Cas9 (see below). SV40-transformed human XP-F fibroblasts XP2YO (XPF-deficient, GM08437), XP2YO complemented with wild-type XPF or mutant XPF-D687A and SV40-transformed human XP-A fibroblasts XP2OS and XP2OS complemented with wild-type XPA were generated by lentiviral transfection as previously described ^23, 42^. SV40-transformed human XP-G fibroblasts XP3BR (XPG-deficient) were from Kaoru Sugasawa (Kobe University) and were complemented with wild-type XPG or XPG-E791A as described previously ^23^. U2OS WT, XPC-KO, CSB-KO, and XPA-KO cells were from Martijn S. Luijsterburg (Leiden University Medical Center) ^43^, and U2OS DDB2-KO and XPF-KO cells were from Hannes Lans, Jurgen A. Marteijn and Wim Vermeulen (Erasmus University Medical Center, Rotterdam) ^44, 45^.

### Cell culture

HAP1 cells were maintained in IMDM containing 4.5 g/L glucose with 10% fetal bovine serum, 1% penicillin-streptomycin, and GlutaMAX^TM^ at 37°C with 5% CO2 (Gibco). XP2YO, XP2OS, and XP3BR cells were maintained in DMEM containing 4.5 g/L glucose and 2mM l-glutamine (Cytiva) with 10% fetal bovine serum (FBS, Millipore) and 1% penicillin-streptomycin (P/S, Gibco) at 37°C with 5% CO2. For survival and COMET chip experiments, U2OS cells were maintained in DMEM containing 4.5 g/L glucose and 2mM l-glutamine (Cytiva) with 10% fetal bovine serum (FBS, Millipore) and 1% penicillin-streptomycin (P/S, Gibco) at 37°C with 5% CO2. For GLOE-seq experiments with U2OS WT, XPC-KO, CSB-KO and XPA-KO cells were maintained in McCoy’s 5a modified cell culture medium (Gibco) at 37°C with 5% CO2.

### Generation of mutant HAP1 cell lines with CRISPR-Cas9

For *XPG-E791A and XPF-D687A HAP1 cell lines:* sgRNAs, single-stranded oligodeoxynucleotides (ssODNs), purified S.p.Cas9 (catalog no. 1081058), and Alt-R HDR Enhancer (catalog no. 1081072) were purchased from IDT. For XPG-E791A knock-in, the following sequences were used; <u>sgRNA</u> (sequence: CACTGCGCCTCTGCTTCCAT): mC*mA* mC*rUrGrCrGrCrCrUrCrUrGrCrUrUrCrCrArUrGrUrUrUrUrArGrArGrCrUrArGrArArArUrA rGrCrArArGrUrUrArArArArUrArArGrGrCrUrArGrUrCrCrGrUrUrArUrCrArArCrUrUrGrArAr ArArArGrUrGrGrCrArCrCrGrArGrUrCrGrGrUrGrCmU*mU*mU*rU, <u>ssODN</u>:/AlT-R HDR1/C*C*TGCGCCTGTTCGGCATTCCCTACATCCAGGCTCCCATGGAAGCGGCCGCGC AGTGCGCCATCCTGGACCTGACTGATCAGACTTC*C*G/AlT-R-HDR2/. For XPF-D68 7A knock-in, the following sequences were used; <u>sgRNA</u> (sequence: CAATGTCAATGC CCCGACGA): mC*mA*mA*rUrGrUrCrArArUrGrCrCrCrCrGrArCrGrArGrUrUrUrUrArGrA rGrCrUrArGrArArArUrArGrCrArArGrUrUrArArArArUrArArGrGrCrUrArGrUrCrCrGrUrUrAr UrCrArArCrUrUrGrArArArArArGrUrGrGrCrArCrCrGrArGrUrCrGrGrUrGrCmU*mU*mU*r U, <u>ssODN</u>: /AlT-R-HDR1/T*G*GCCAGGAACAGAATGGTACACAGCAAAGCATAGTTG TGGCAATGCGTGAATTTCGAAGTGAGCTTCCATCTCTGATCCATCGTCGGGGCATT GACATTGAACCCGTGACTTTAGAGGTTG*G*A/AlT-R-HDR2/. CRISPR ribonucleoprotein (RNP) complexes were transfected into HAP1 cells using CRISPRMAX^TM^ (Thermo F isher Scientific). 0.5 μL of sgRNA (3 μM), S.p.Cas9 (3 μM), and Cas9 Plus Reagent (0.5 μL) were mixed with 10.5 μL Opti-MEM^TM^ (Thermo Fisher Scientific) and incubated at room temperature (RT) for 10 minutes. 120 nM of RNP (12.5 μL) was mixed with 0.3 μL C RISPRMAX^TM^ Reagent and 5 μL Opti-MEM^TM^ and incubated at RT for 10 minutes. The RNP/liposome complex was added into a well of 96-well plate with 0.4 × 10^4^ cells in tota l IMDM growth medium (125 μL), followed by addition of 1.25 μL 3 mM Alt-R HDR Enha ncer. Cells with RNP complex were incubated at 37°C with 5% CO2 for 72 hours. The m utant genome loci were amplified with PCR primers to validate the mutation (XPG: Forw ard-CCCTGGGGGAATGCACTGCATG, Reverse-CCACAAGCTTCTGCCTCAGCCC; X PF: Forward-CCCAGCTCCTTCCCTTTCCCCA, Reverse-ACAACTCCGCCGTTGCATG AGG). Sanger sequencing was performed to determine the sequence of mutant alleles; XPG-E791A: AGAG(E) → GGCC(A) and XPF-D687A (additional R deletion at 701): GA T(D) → GCA(A) and TCG (R) deletion.

For *ERCC1 knock out HAP1 cell lines:* Exon 4, which encodes part of the XPA binding domain, was disrupted with a hygromycin B resistance marker. Two gRNA sequences targeting exon 4 (TTGCGCACGAACTTCAGTAC and AATTACGTCGCCAAATTCCC) were cloned into pX330 vector. Sequences upstream and downstream of exon 4 were amplified with the following primers: GCCAAGCCCTTATTCCGATCTACAC and GAAGGGCAGAAGCCATCAATAGGG for the left arm, GTGAGCTCTGCGGCGCCACC and GGAATACTAAGGGCTCAGAGTACGGC for the right arm. These left arm and right arm sequences were then cloned into the targeting vector DT-A-pA/loxP/PGK-Hygro-pA/loxP (Laboratory for Animal Resources and Genetic Engineering, Center for Developmental Biology, RIKEN Kobe, gift from Professor S. Takeda) flanking the hygromycin resistance gene. gRNA and targeting vectors were transfected into HAP1 cells (C859, Horizon) using Xfect Transfection Reagent (Takara). Integration of the hygromycin resistance marker in exon 4 was also confirmed by PCR with the following primers ATCTTTGTAGAAACCATCGGCGCAGCTATT (anneals to hygromycin resistance gene) GGGAGTTGAGAGGTCTCAGTCTCTTC (anneals to ERCC1 gene sequence downstream of the right arm).

### Trabectedin or UV irradiation for alkaline COMET chip assay

Cells were enriched at the G1 phase by incubating in growth medium supplemented with 1 μM (U2OS) or 2 μM (HAP1) palbociclib (hereafter referred to as the “working medium”) for 24 hours prior to exposure to trabectedin or UV irradiation. XP2YO, and XP2OS were asynchronous prior to exposure to trabectedin or UV irradiation. Following this, cells were embedded in a COMET chip and further incubated for 30 minutes at 37°C in the working medium. The medium was removed, and the cells were incubated in the working medium supplemented with 50 nM trabectedin for 2 hours in the presence and absence of repair synthesis inhibitors (HAP1: 0.5 mM HU, 5 μM AraC, U2OS: 1 mM HU, 10 μM AraC, XP2YO and XP2OS: 4 mM HU, 40 μM AraC). After trabectedin treatment, cells were incubated in the working medium either with or without the repair synthesis inhibitors for varying repair periods. In the case of UV treatment, after removal of the medium, cells were subjected to irradiation using 5 J/m^2^ of UV-C (254 nm UV light). This was followed by incubation at 37°C in the working medium, with or without the repair synthesis inhibitors, for different repair times. DNA strand breaks were examined utilizing the alkaline COMET chip assay.

### Alkaline COMET chip assay

The high-throughput variant alkaline COMET chip assay was performed as previously 22,46, with modification in the unwinding and electrophoresis steps. Unwinding of DNA was carried out for 30 minutes twice (HAP1, U2OS); 15 minutes twice (XP2YO, XP2OS). Electrophoresis was carried out for 50 minutes, 1 V/cm at 4°C (HAP1, U2OS); 30 minutes, 1 V/cm at 4°C (XP2YO, XP2OS). Both unwinding and electrophoresis steps were conducted in alkaline electrophoresis buffer (200 mM NaOH/1 mM EDTA/0.1% Triton X-100). 30 μm CometChip^®^ (catalog no. 4250-096-01), low melting agarose (catalog no. 4250-500-02), and lysis solution (catalog no. 4250-500-01) were from Trevigen. 1X SYBR Gold (Invitrogen) was used for staining and comets were imaged with 4X magnification on a fluorescence microscope (BX53, Olympus). % DNA in tail was quantified with Comet analysis software (catalog no. 4260-000-CS) from Trevigen.

### Clonogenic survival assay

HAP1, U2OS, XP2YO, XP2OS, and XP3BR cells were cultured in growth media (refer to the Cell culture section). 1500 cells were seeded in triplicated 6 cm dishes a day before trabectedin or illudin S treatment. Cells were treated for 2 hours with growth media containing either trabectedin or illudin S in varied concentration. Following treatment, media were changed to fresh growth media and cells were grown for 7-8 days. Cells were fixed with 4% paraformaldehyde for 15 minutes and stained with 1% methylene blue for 2hours. After washing with water, colonies (defined as ≥ 25 cells) were counted. The survival rate was normalized to the number of colonies of non-treated cells.

### Cell lysis and Western blotting

Cells were rinsed with ice-cold PBS and lysed in M-PER buffer (Thermo Fisher Scientific) containing Halt protease and phosphatase inhibitor cocktail (1X) (Thermo Fisher Scientific). The concentration of protein was determined using a Bio-Rad DC Protein Assay Kit (Bio-Rad). Samples were prepared by adding LDS sample buffer containing 2.5% of 2-mercaptoethanol (4X) (Invitrogen), followed by boiling at 95 °C for 5 minutes. Samples containing 25 µg of protein were resolved on 8-16% Tris-Glycine or 4-12% Bis-Tris gels (Thermo Fisher Scientific) at 150V for 45 minutes, transferred onto Amersham Hybond 0.2 μm PVDF membrane (GE Healthcare) at 250 mA for 70 minutes with mini-protein tetra system (Bio-Rad). Transferred membranes were blocked with 5% skim milk in Tris-buffered saline (20 mM Tris Base, 137 mM NaCl, pH 7.6) containing 0.1% Tween 20 (TBS-T) for 1 hour at room temperature (RT) and incubated with the following antibodies : α-β-actin (mouse, 1:10,000), α-XPG (rabbit, 1:500), α-XPF (rabbit, 1:2,000), or α-ERCC1 (mouse, 1:300) were added to TBST and incubated overnight at 4°C. Blots were then incubated with anti-goat IgG rabbit or mouse antibody (1:2,000) for 1 hour at RT. Detection of horseradish peroxidase-conjugated secondary antibodies was performed with SuperSignal West Pico PLUS Chemiluminescent Substrate (Thermo Fisher Scientific) under ChemiDoc touch imaging system (Bio-Rad).

### Analysis of survival and COMET assay data

Quantitative values were expressed as mean ± SEM or mean ± SD. The exact sample size for each experiment is described in figure legends. Statistical significance of the survival and COMET assay data were analyzed by performing the ordinary two-way ANOVA or two-tailed paired t-tests. Differences between groups were considered significant when P<0.05. These statistical analyses were performed using GraphPad Prism (GraphPad Software) or Microsoft Excel.

### GLOE-seq library preparation

For the GLOE-seq positive control, *i.e.*, checking the method’s ability to identify DNA breaks introduced at known genomic locations (**Supplementary Fig. 4a-b**), genomic DNA was extracted using the Monarch genomic DNA purification kit (T3010) from untreated U2OS wildtype cells after harvesting (5 min incubation with 0.25 % trypsin). Existing breaks were blocked by incubating 5 μg DNA first with 10 units of T4 PNK per 4 ng DNA and 1x NEBuffer 2 for 30 min at 37 °C, and then with 250 μM ddNTP mix, 2 units of Therminator IX DNA polymerase and 1x ThermoPol buffer for 10 min at 60 °C. DNA was purified with ProNex size-purification system (NG2001, Promega) with 8:5 beads to sample ratio. DNA was incubated with 2.5 units of Nb.BsrDI and 1x CutSmart buffer for 90 min at 65 °C and then with 5 units antarctic phosphatase and 1x antarctic phosphatase buffer for 30 min at 37 °C, and purified as described above. DNA library for Illumina sequencing was prepared according to GLOE-Seq protocol ‘Steps y23 – 28: Denaturation and ligation of 3’-OH termini (yeast)’ onwards ^28^.

In the rest of GLOE-seq experiments, cells were grown up to 60-80 % confluency and incubated for 24 h with 1 mM palbociclib isethionate for G1 enrichment. Cells were then treated with 0.1 % DMSO, 20 nM or 50 nM trabectedin in combination with 1mM palbociclib isethionate for 2 h. Cells were washed with PBS and underwent a 2-h recovery in cell culture media without trabectedin in the presence of 1 mM palbociclib isethionate. Nuclei isolation and DNA library preparation for sequencing followed the GLOE-seq protocol for mammalian cells ^28^ with the following modifications. *Isolation of genomic DNA from mammalian cell culture:* Agarose plugs were incubated with 6 mL of proteinase K solution and shaken at 170 rpm*. Fragmentation and capture of biotinylated single-stranded DNA:* DNA was sonicated with an average fragment length of 300 nucleotides with Qsonica for 5 min at 4 °C in cycles of 15 s ON and 5 s OFF with 20 % amplitude. DNA was purified twice with AMPure beads with elution in 50 μL dH2O. Streptavidin MyOne C1 dynabeads were resuspended after washing in 50 μL bind and wash buffer (2x) for a 1:1 beads to sample ratio. *Second strand synthesis, end polishing and ligation of the distal adaptor:* Reagents for second strand synthesis, end polishing and distal adaptor ligation were combined on ice and mixed by pipetting. For the distal adaptor either 3792-UMI or 3792-UMI.v2 (different indexes) were annealed with 3791. DNA was purified with AMPure beads and eluted in 30 μL of dH2O. *qPCR:* Before the final library amplification described in ^28^, qPCR with reaction conditions identical to the final PCR used for library amplification was done for quality control. Samples (1 μL) were diluted in deionized water (9.4 μL) and combined with 1x Q5 buffer, 0.2mM dNTPs, 0.1 μM i5 illumina indexing primer, 0.1 μM P7-short primer, 1x EvaGreen and 0.1 units of Q5 high-fidelity DNA polymerase. qPCR protocol included 120 s initial denaturation at 95 °C, 40 cycles of 15 s denaturation at 95 °C, 30 s annealing at 60 °C and 20 s extending at 72 °C, as well as 95 s final extension at 72 °C. *Library amplification:* The library was scaled up from 20 μL to 50 μL using entire samples. A customized P7-short primer compatible with the UMI-containing distal adaptors was used. Samples were purified with AMPure beads with a 1:1 bead to sample ratio. The DNA libraries were sequenced on an Illumina NovaSeq 6000 with a single-read protocol and the read length of 100 bp (R1); additionally, 15 cycles were run on R2 to read UMI and custom indexes introduced by the oligos 3792-UMI or 3792-UMI.v2. Supplementary Table 1 summarizes the reagents, enzymes, kits, oligonucleotides, and other materials used for GLOE-seq experiments.

### GLOE-seq data analysis

#### Sequencing read processing

After demultiplexing of sequencing data, each sample was represented by two fastq.gz files, with the first file containing 101-nucleotide-long genomic reads (R1) and the second storing the respective 10-nucleotide long UMIs (R2). The quality of the R1 and R2 data was checked using FastQC/0.11.9 ^47^. Low-quality and adaptor-containing reads were removed via trimmomatic/0.38 ^48^, using the following parameters: for R1, SE ILLUMINACLIP:Trimmomatic-0.39/adapters/TruSeq3-SE.fa:2:30:10 LEADING:3 TRAILING:3 SLIDINGWINDOW:4:15 MINLEN:101; for R2, SLIDINGWINDOW:4:15 MINLEN:10. Using a custom Python/3.7.4 script employing the module Biopython/1.79, we merged the R1 and R2 records that passed trimming, incorporating the 10-nucleotide-long R2 sequences in the name of respective R1 reads. These reads were mapped to human reference genome GRCh38 via bowtie2/2.3.5.1, using the pre-built bowtie2 ^49^ index from https://genome-idx.s3.amazonaws.com/bt/GRCh38_noalt_as.zip and default settings. Read duplicates were removed by the tool dedup of umi_tools/1.1.2 toolkit ^50^, grouping reads with the same R2 sequence stored in the read name (*method=unique*). samtools/1.12 ^51^ were employed to remove unmapped reads, sort, index and generate statistics of bam files. bedtools2/2.29.2 ^52^ was used to covert bam files to bed files. Each read represented one unit of DNA-break signal, which we positioned at the nucleotide located immediately upstream of the 5’ end of the read. Since a GLOE-Seq read is the reverse complement of the DNA fragment captured in the protocol ^28^, the strand of the nucleotide bearing the signal was changed to the opposite to the one on which the respective read was mapped. In this way signal positioning, the original DNA break is on the 5’ side of the nucleotide bearing the signal. Using AWK and a custom Python/3.7.4 script with the modules numpy/1.21.5 and pandas/0.25.1, we implemented the described DNA-break-signal positioning, which resulted in sample-specific BED files containing the genomic coordinates of the nucleotides bearing the signal (immediately downstream of a break), the DNA-break scores (always one) and MAPQ scores.

#### Software for mapped DNA break data analysis

The downstream analysis of DNA break data and their visualization were performed via custom Python/3.7.4 scripts and Jupyter notebooks employing the modules numpy/1.19.2, scipy/1.6.3, pandas/1.1.3, biopython/1.79, matplotlib/3.4.2 and seaborn/0.11.1 in Python/3.8.5 environment. Besides, bedtools2/2.29.2 ^52^ was used to extract the sequence context of DNA breaks from the reference genome (GRCh38). The custom code for genome-scale data analysis is available at https://gitlab.ethz.ch/eth_toxlab/trabi-seq.

#### External datasets

The following publicly available datasets were used in the analysis. Human reference genome, GRCh38: pre-built bowtie2 index from https://genome-idx.s3.amazonaws.com/bt/GRCh38_noalt_as.zip. Transcript coordinates: GENCODE/V41/knownGene, retrieved from UCSC Table Browser. Canonical transcripts of genes: GENCODE/V41/knownCanonical, retrieved from UCSC Table Browser. Gene expression: CCLE_expression_full from DepMap Public 22Q2 (https://depmap.org/portal/download/all/), with the cell-line accession numbers ACH-000364 (U2OS WT) and ACH-002475 (HAP1 WT). Protein-coding genes: GENCODE/V41/knownToNextProt, retrieved from UCSC Table Browser. Coordinates of centromeres and gaps: retrieved from UCSC Table Browser for GRCh38. Chromatin accessibility and histone modification: from GSE87831 (GEO) ^53^, GSE87831_DNase-Seq.r1.peaks.bed.gz, GSE87831_DNase-Seq.r2.peaks.bed.gz, GSE87831_H3K4me3.peaks.bed.gz (referred to as H3K4me3 [1] in **Fig. 3c**); from GSE44672 (GEO) ^54^, GSM1356566_U2OS_H3K4me3.txt.gz (referred to as H3K4me3 [2] in **Fig. 3c**), GSM1356565_U2OS_H3K4me1.txt.gz, GSM1356567_U2OS_H3K27ac.txt.gz ; genomic coordinates were converted from hg19 to hg38 via https://genome.ucsc.edu/cgi-bin/hgLiftOver. Oncogenes: COSMIC Cancer Gene Census (downloaded on 15.05.2023).

#### Gene boundaries, DNA break count normalization and further figure details

In our analysis, genes were represented by canonical transcripts (GENCODE/V41/knownCanonical), whose boundaries were considered as transcription start site (TSS) and transcription end site (TES). When operating with whole-gene features, we summed the signals mapped between TSS and TES. Genome-scale data required normalization of mapped signals since sequencing depth varied across samples. Therefore, for each sample, we calculated a normalization factor that reflects the level of endogenous DNA breaks in unexpressed genes. Specifically, the normalization factor was 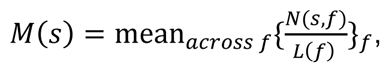 where features *f* here are the transcribed (antisense) strands of not expressed genes, *N*(*s*, *f*) is the number of mapped DNA breaks per a concrete feature *f* in the sample *s*, and *L*(*f*) is the feature’s length measured in kilobases. DNA break count data *C*(*s*, *f*) were related to these sample-specific normalization factors via the formula 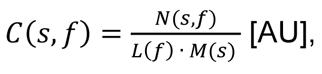, where a feature *f* is a chromosome bin with both strands considered together (**Fig. 3b**), a gene’s transcribed strand or a gene’s non-transcribed strand (**Fig. 3d-g**, **Supplementary Fig. 5a-f**), 5-kilobase region downstream or upstream of the TSS with both strands considered together (**Fig. 4g-h, Supplementary Fig. 7**). In **Supplementary Fig. 4c**, DNA break counts per bin were further related to the median value in a sample (fold change with respect to the median,

*C*(*s*, *f*)/median_*acrossss* *f*_{*C*(*s*, *f*)}_*f*_). In **Fig. 3h-k** and **Supplementary Fig. 5g**, we presented mean_*acrossss* *f*_{*C*(*s*, *f*)}_*f*_.

For profiles of mean DNA break counts (**Fig. 4a-f, Supplementary Fig. 6a-f**), the following data were presented: 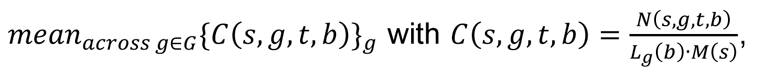 where *N*(*s, g, t, b*) is the number of mapped DNA breaks per bin *b* in sample *s*, gene *g* and strand *t*, *L*_g_*(b)* is the bin’s length measured in kilobases, *M*(*s*) is the normalization factor, and *G* is the set of top 30% expressed protein-coding genes in unexposed U2OS WT. When a bin size was absolute (provided as a base number), bins had the same size across all considered genes, *L*_*g*_(*b*) = *L*(*b*). When a bin size was relative (provided as a percentage, specifically, in the region between TSS and TES), each bin was the indicated fraction (*α*) of the respective gene length, therefore, the bin size *L*_*g*_(*b*) = *a* · *L*(*g*) measured in bases varied across genes (the average value is indicated in parentheses in respective figure panels).

In U2OS, we considered 16,740 protein-coding genes, including not expressed genes: 1,989, ≤ 25% gene expression tier: 3,689, ≤ 50%: 3,689, ≤ 70%: 2,948, ≤ 80%: 1,475, ≤ 90%: 1,475, ≤ 95%: 737, ≤ 100%: 738. Number of genes beyond the maximal Y-axis value: in Fig. 3d, 16 (TS, transcribed strand), 0 (NTS, non-transcribed strand); in Fig. 3e, 0 (TS), 1 (NTS); in Fig. 3f, 34 (TS), 0 (NTS); in Fig. 3g and Supplementary Fig. 5a,c,d, 0 (TS), 0 (NTS); in Supplementary Fig. 5b, 0 (TS), 2 (NTS). In HAP1, we considered 16,740 protein-coding genes, including not expressed genes: 3,428, ≤ 25% gene expression tier: 3,334, ≤ 50%: 3,324, ≤ 70%: 2,660, ≤ 80%: 1,331, ≤ 90%: 1,331, ≤ 95%: 666, ≤ 100%: 666. Number of genes beyond the maximal Y-axis value: in Supplementary Fig. 5e, 19 (TS), 26 (NTS); in Supplementary Fig. 5f, 10 (TS), 3 (NTS).

## DATA AVAILABILITY

Raw sequencing data and BED files with called DNA breaks are deposited at the NCBI Gene Expression Omnibus (GEO) (https://www.ncbi.nlm.nih.gov/geo/); accession number pending (once generated, the accession number will be provided on the gitlab page indicated below). Supplementary Table 1 and read statistics are available along with the code at https://gitlab.ethz.ch/eth_toxlab/trabi-seq.

## CODE AVAILABILITY

The code for genome-scale data analysis required to generate respective figures is available at https://gitlab.ethz.ch/eth_toxlab/trabi-seq.

## ACKNOWLEDGEMENTS

Genome-scale data produced and analyzed in this paper were generated using the resources of the ETH Zürich Genetic Diversity Centre (GDC), the Functional Genomics Center Zürich (FGCZ), and the ETH Zürich Euler cluster. The authors thank members of Laboratory of Toxicology at ETH Zürich and the SNF Sinergia consortium “Hijacking Transcription-Coupled DNA Repair for Cancer Therapy” for productive discussion of the work. The authors are grateful to Martijn Luijsterburg (University of Leiden), and Hannes Lans, Jurgen Marteijn and Wim Vermeulen (Erasmus MC) for U2OS cell lines, Jiyoung Park (IBS-CGI) for help with establishing COMET chip assay, Navnit Kaur Singh, Laura Slappendel, Jasmina Büchel and Sabine Diedrich for help with experiments, and Sharon Cantor (UMass Worcester) for a critical reading of the manuscript. This work was supported by the Korean Institute of Basic Science (IBS-R022-A1 to ODS, IBS-R022-A2 to DI), the Swiss National Science Foundation (Sinergia grant CRSII5-186332 to SJS and ODS) and the Deutsche Forschungsgemeinschaft (DFG, German Research Foundation – Project-ID 393547839 – SFB 1361 to HDU).

## AUTHOR CONTRIBUTIONS

KS, VT, SJS, and ODS conceptualized research. KS, VM, and HY conducted COMET chip and survival assays. KS, HY, and DI generated cell lines. VT designed experiments for genome-scale DNA break mapping and analyzed the respective data. ED prepared GLOE-seq libraries. NJLP helped to implement the GLOE-seq protocol. NZ and HDU upgraded the GLOE-seq protocol and advised on its implementation. KS, VT, and ODS wrote the original manuscript draft, and it was reviewed by all the authors. ODS, SJS and HDU supervised research and acquired funding.

## COMPETING INTERESTS

The authors declare that they have no conflict of interest.

---

**Supplemental Fig. 1:**
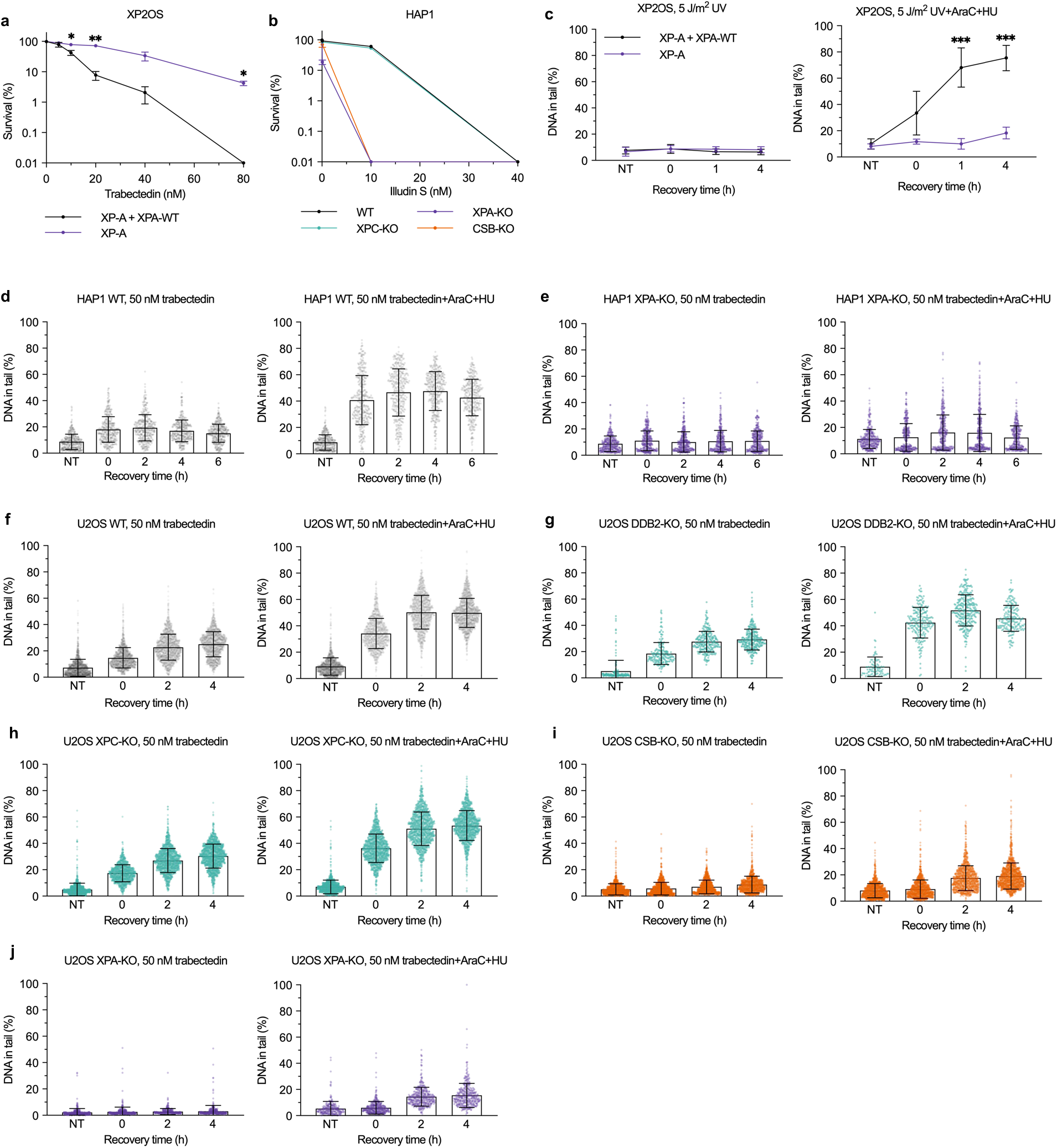
Related to Fig. 1. **a**, XP2OS (XP-A or lentiviral expression of XPA-WT) cells were treated with the indicated doses of trabectedin or DMSO control for 2 h and incubated with fresh medium. The number of colonies was counted after 8 days. Mean ± SEM of 3 biological replicates (3 technical replicates per experiment). *P<0.05, **P<0.01 using two-tailed paired t-test (between XP-A and XP-A+XPA-WT at each concentration). **b**, HAP1 WT, XPC-, XPA-, and CSB-KO were treated with 0.1, 10, and 40 nM illudin S or DMSO control for 2hrs and incubated with fresh medium. The number of colonies was counted after 8 days. Mean ± SEM of 2 biological replicates (3 technical replicates per experiment). **c**, XP2OS cells were treated trabectedin (50 nM, 2 h) and incubated for 1 or 4 h with or without repair synthesis inhibitors (4 mM HU, 40 μM AraC). ssDNA breaks were analyzed by alkaline COMET chip assays. Mean ± SEM of 3 biological replicates. ***P<0.001 using ordinary two-way ANOVA with uncorrected Fisher’s LSD (between XP-A and XP-A+XPA-WT at each recovery time). **d-j**, DNA in tail (%) from all comets used for Fig. 1d and e (a dot represents DNA in tail (%) of a comet analyzed; error bars represent SD).

**Supplementary Fig. 2.**
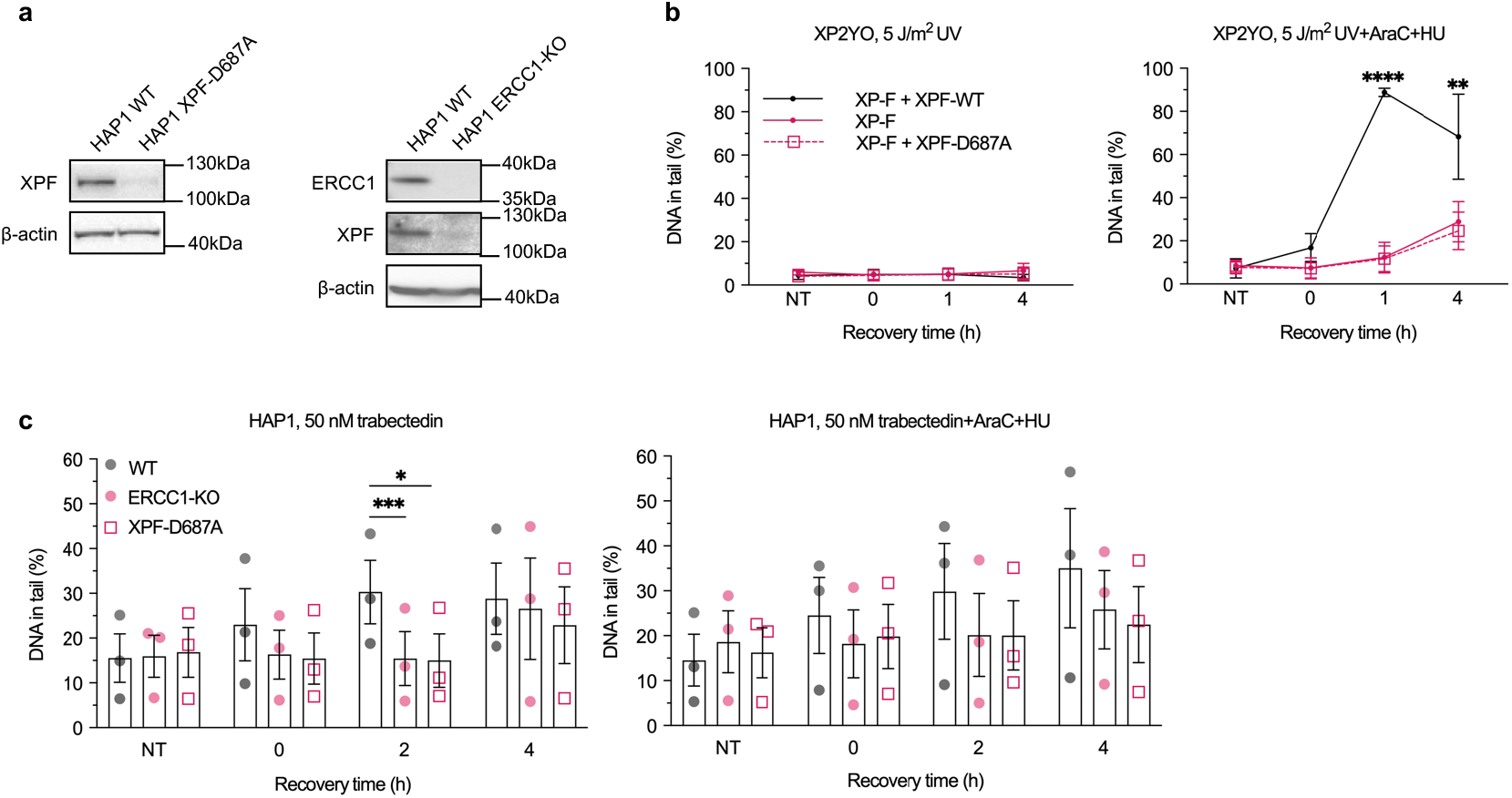
Related to Fig. 2. **a**, Cells were lysed and subjected to Western blot analysis for XPF and ERCC1 expression (n=3). **b**, XP2YO patient cells (XP-F, with or without lentiviral expression of XPF-WT or XPF-D687A) were exposed to 5 J/m^2^ UV-C and allowed to repair for 1 or 4 h with or without repair synthesis inhibitors (4 mM HU, 40 μM AraC). ssDNA breaks were analyzed by alkaline COMETchip assays. Mean ± SEM of 3 biological replicates. **P<0.01, ****P<0.0001 using ordinary two-way ANOVA with Dunnett’s multiple comparisons test (between XP-F+XPF-WT and XP-F or XP-F+XPF-D687A at each recovery time). **c**, Fig.2b in dot plots. *P<0.05, ***P<0.001 using two-tailed paired t-test (between WT and ERCC1-KO or XPF-D687A at each recovery time).

**Supplementary Fig. 3.**
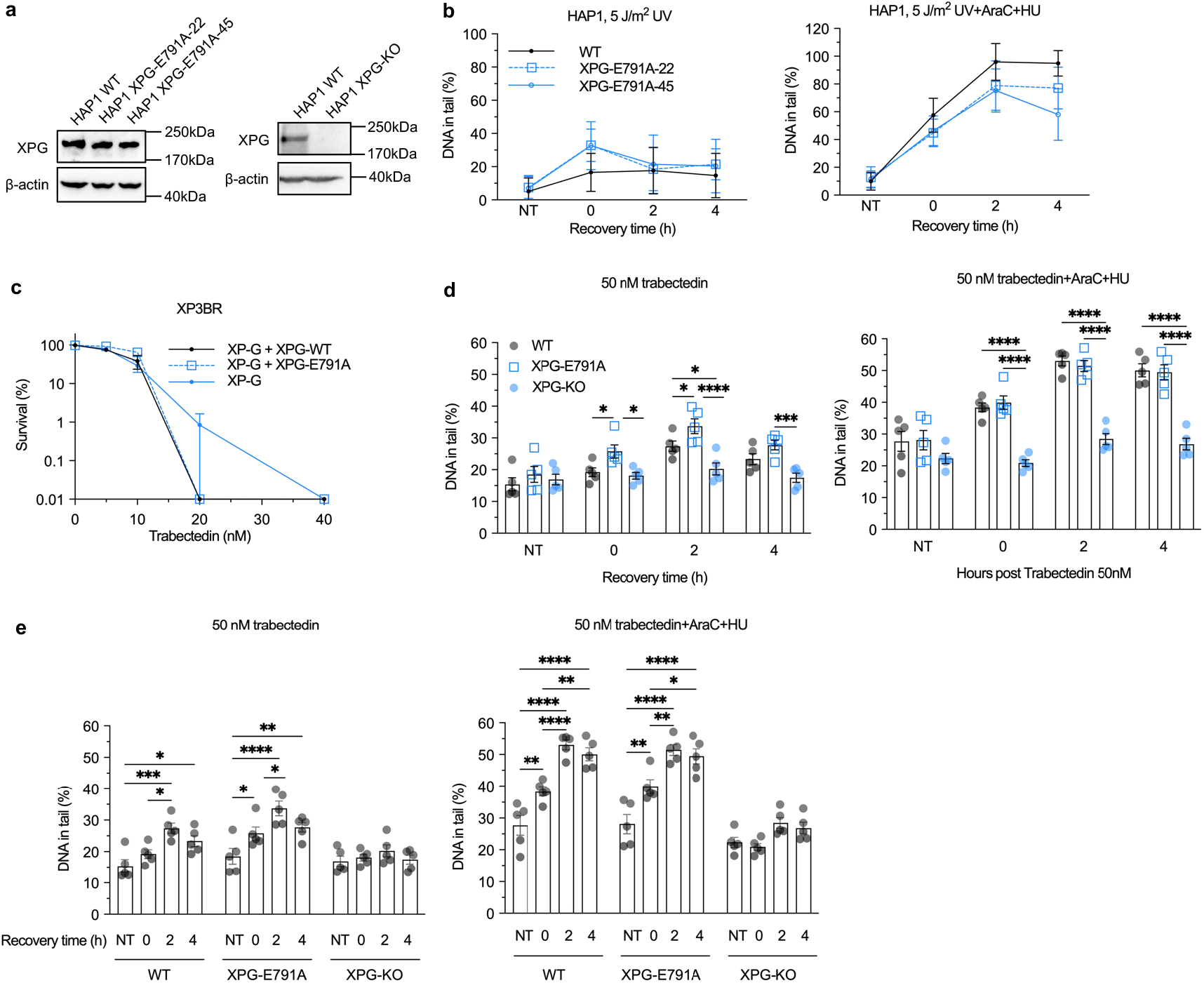
Related to Fig. 2. **a**, Cells were lysed and subjected to Western blot analysis for XPG expression (n=3). **b**, HAP1 WT, XPG-E791A clones #22 and #45 were arrested in G1 with palbociclib (2 µM, 24 h), exposed to 5 J/m^2^ UV-C and allowed to repair for up to 4 h with or without repair synthesis inhibitors (0.5 mM HU, 5 μM AraC). ssDNA breaks were analyzed by alkaline COMET chip assay (n=1). Error bars represent the polydispersity of the individual comets. **c**, XP3BR patient cells (XP-G, with or without lentiviral expression of XPG-WT or XPG-E791A) were treated with the indicated doses of trabectedin or DMSO control for 2 h and incubated with fresh medium. The number of colonies was counted after 8 days. Mean ± SEM of 2 biological replicates (3 technical replicates per experiment). **d-e**, Fig. 2f in dot plots. *P<0.05, **P<0.01, ***P<0.001, ****P<0.0001 using ordinary two-way ANOVA with Tukey’s multiple comparisons test. Comparison of WT, XPG-E791A, and XPG-KO at each recovery time (**d**); and between the recovery time for the individual cell lines (**e**).

**Supplementary Fig. 4.**
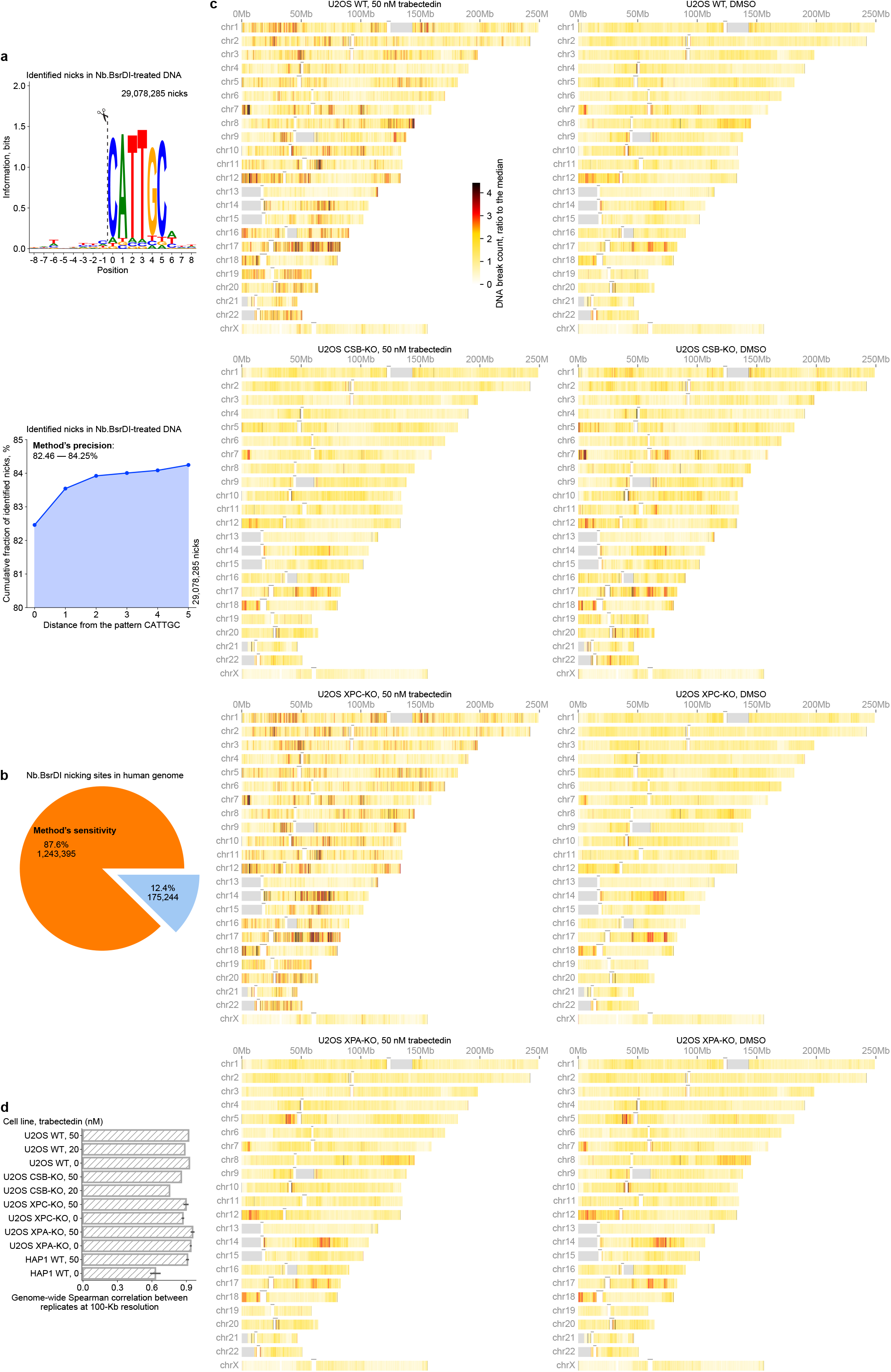
Related to Fig. 3. Positive control of GLOE-Seq and genome-wide distribution of DNA breaks. **(a)** Upper panel: Sequence logo showing sequence conservation around the DNA breaks (dashed line) identified in naked DNA that, first, underwent blocking of existing (i.e., endogenous and handling-associated) breaks and, second, was treated with the nicking endonuclease Nb.BsrDI. Lower panel: Cumulative fraction of identified DNA breaks along the increasing distance from the 5’ end of the Nb.BsrDI recognition pattern CATTGC. The non-zero distance from the 5’ end of Nb.BsrDI recognition pattern corresponds to potential sloppiness (imperfect specificity) of the endonuclease. Precision, *i.e.*, the fraction of DNA breaks introduced by Nb.BsrDI, was estimated considering the distances 0 and 5 (82.46-84.25%). **(b)** Sensitivity estimated as the fraction of detected Nb.BsrDI recognition patterns CATTGC in the human reference genome. Assuming the endonuclease imperfect specificity and the distance smaller or equal to 5 (a), the sensitivity is 87.65%, whereas it is 87.32% when only the distance 0 is considered. The method can identify multiple breaks at one location (**Fig. 3a**), therefore, the number of identified DNA breaks in **a** is higher (around 20-fold) than the number of recognition patterns in the genome. **(c)** Genome-wide distribution of DNA breaks at 100-Kb resolution across cell lines with and without trabectedin exposure. Presented data: one biological replicate per cell line. Gray areas correspond to heterochromatin and short arm gaps in the human reference genome. Horizontal black dash above each chromosome indicates centromeric regions, for which data are not shown. Read number normalization is described in **Methods**. DNA break count values are capped at 4.44. **(d)** Reproducibility of genome-wide DNA break distribution at 100-Kb resolution across biological replicate experiments. Bars and errors: mean ± s.d. of Spearman correlation coefficient across pairs of biological replicates (no errors: cases when only two replicates are available).

**Supplementary Fig. 5.**
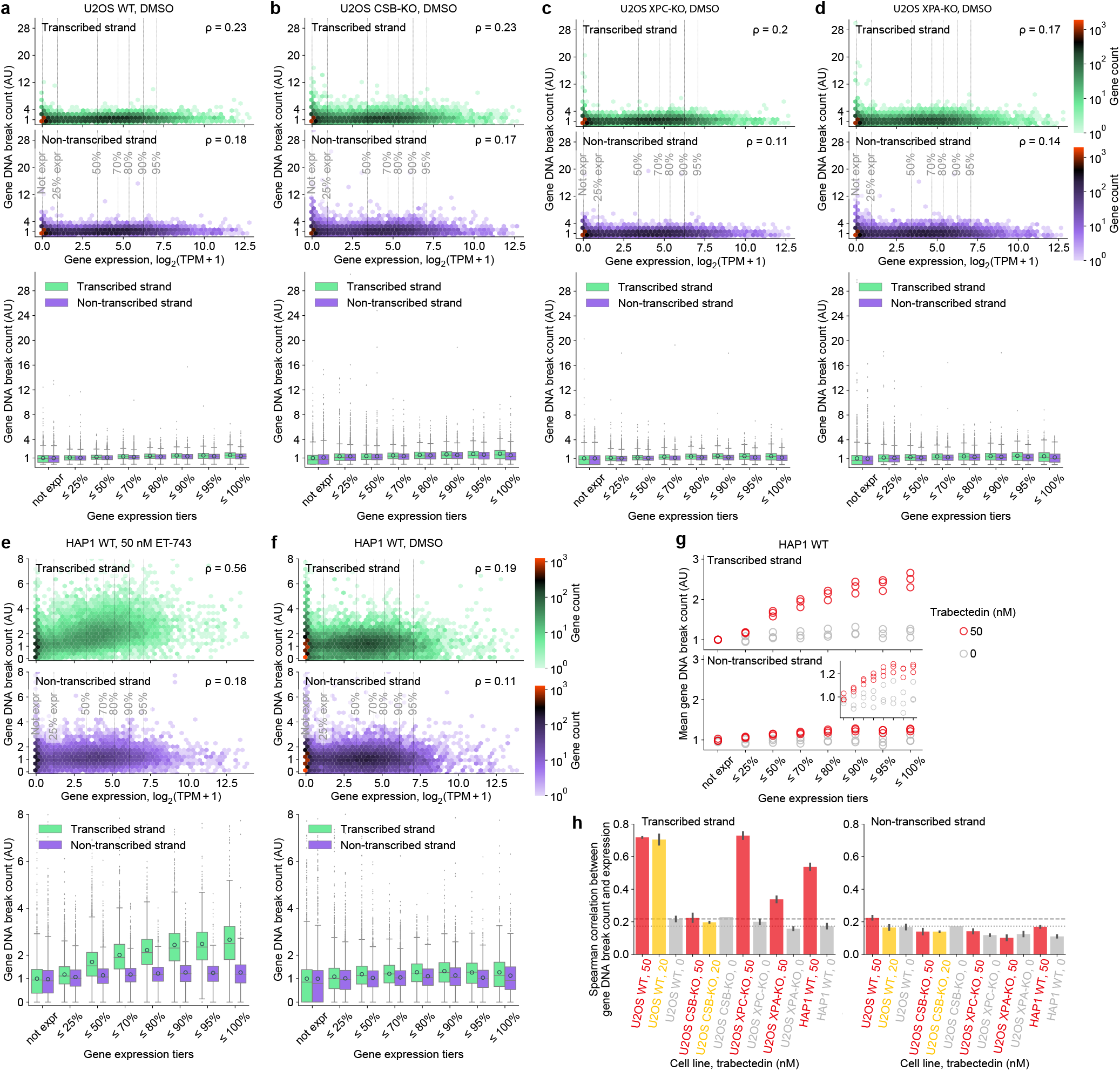
Related to Fig. 3. **(a-d)** DNA break count on the transcribed and non-transcribed strands of protein-coding genes in U2OS WT (a), CSB-KO (b), XPC-KO (c) and XPA-KO (d) after two-hour exposure to DMSO and subsequent two-hour recovery versus gene expression level in unexposed U2OS WT. **(e-f)** DNA break count on the transcribed and non-transcribed strands of protein-coding genes in HAP1 WT after two-hour exposure to trabectedin **(e)** or DMSO **(f)** and subsequent two-hour recovery versus gene expression level in unexposed HAP1 WT. **a-f**: The plots are built analogously to Fig. 3d-g. **(g)** Mean DNA break count of protein-coding genes in HAP1 WT after two-hour exposure to trabectedin or DMSO and subsequent two-hour recovery versus gene expression level in unexposed HAP1 WT. The plot is built analogously to Fig. 3h-k. **(h)** Correlation between gene break count on an indicated strand and gene expression across cell lines and trabectedin exposure concentrations. Bar and error: mean ± s.d. across biological replicates. Horizontal lines: mean correlation values for the transcribed strand in unexposed U2OS WT and HAP1 WT; these lines highlight that without chemical exposure, DNA break count correlates with gene expression stronger for the transcribed strand than for the non-transcribed one.

**Supplementary Fig. 6.**
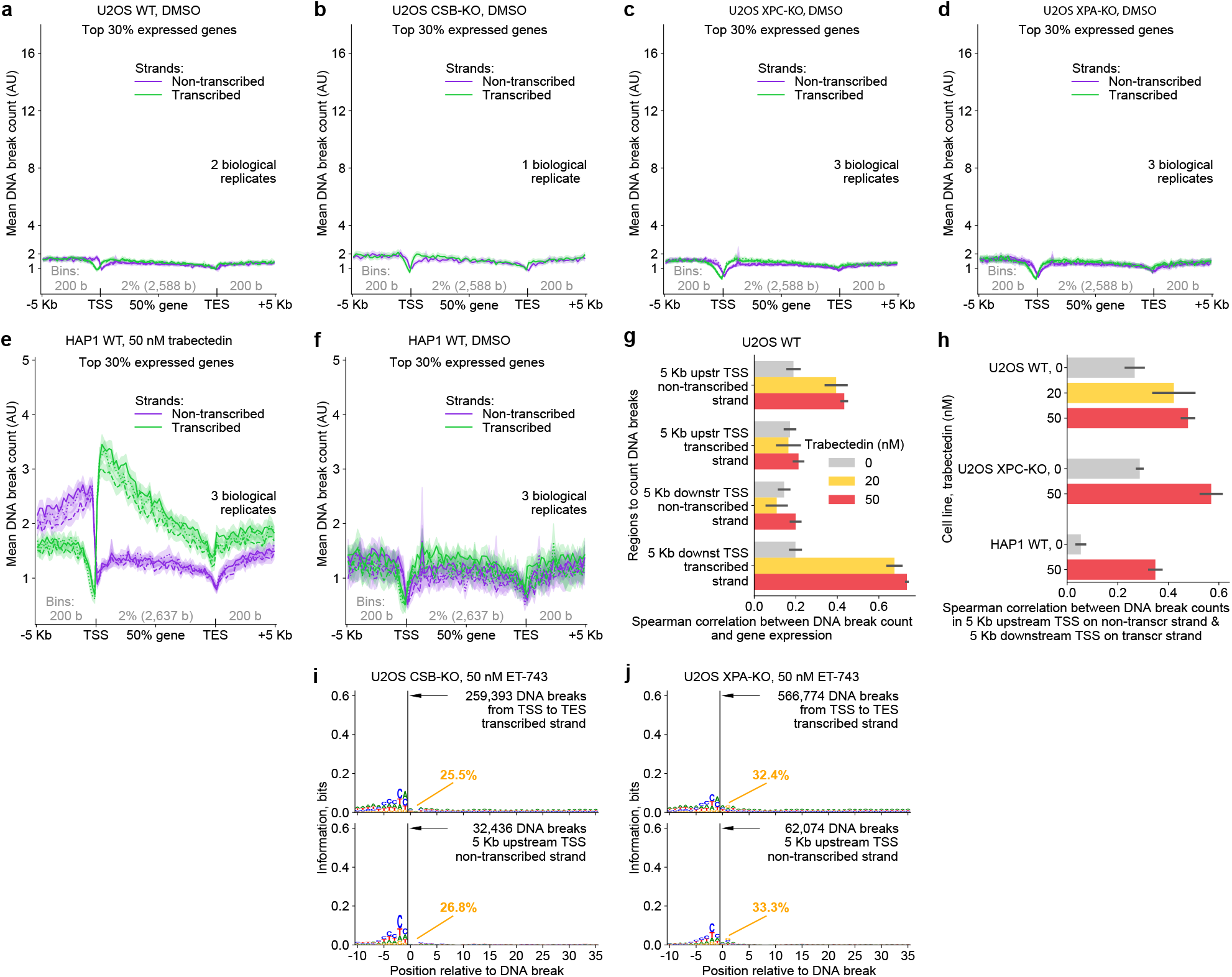
Related to Fig. 4. **(a-d)** Strand-specific profile of the mean DNA break count throughout the gene body and its upstream and downstream 5 kilobase regions in U2OS WT (a), CSB-KO (b), XPC-KO (c) and XPA-KO (d) after two-hour exposure to DMSO and subsequent two-hour recovery. **(e-f)** Strand-specific profile of the mean DNA break count throughout the gene body and its upstream and downstream 5 kilobase regions in HAP WT after two-hour exposure to trabectedin (e) or DMSO (f) and subsequent two-hour recovery. 3,994 protein-coding genes, specifically, top 30% expressed in unexposed U2OS WT, are considered. a-f: The plots are built analogously to Fig. 4a-d. **(g)** Correlation between DNA break count in the indicated regions and gene expression in U2OS WT exposed to 0, 20 and 50 nM trabectedin for two hours with subsequent two-hour recovery. **(h)** Correlation between DNA break counts along two branches of divergent transcription in TC-NER proficient cell lines. g-h: all protein-coding genes are considered. **(i-j)** Sequence logos showing no sequence enrichment around DNA breaks in U2OS CSB-KO (i) and XPA-KO (j). The percentage of G at position 1 (+2 relative to the break) is shown. The plots are built analogously to Fig. 4i-j.

**Supplementary Fig. 7.**
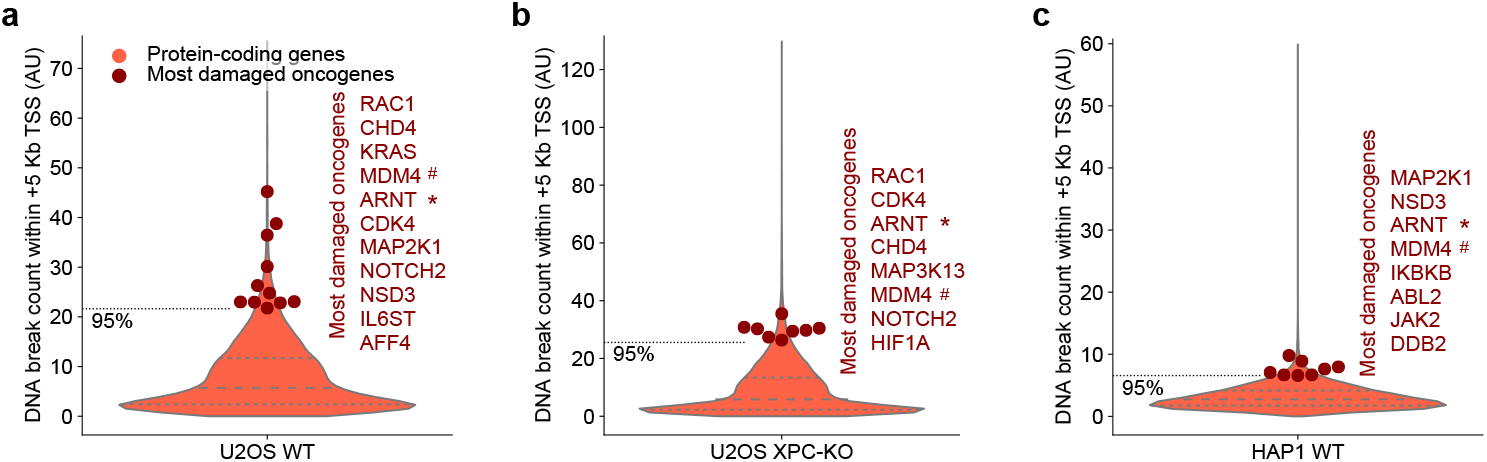
Related to Fig. 4. **(a-c)** The distribution of gene DNA break counts in U2OS WT (a), U2OS XPC-KO (b) and HAP1 WT (c) across protein-coding genes (16,740). Most damaged oncogenes (higher than 95^th^ percentile of the total distribution) are shown as markers and named in the order of descending DNA break count. #, *: the oncogenes found in the group of the most damaged genes across the three cell lines. Oncogene annotation: COSMIC Cancer Gene Census, tier 1, genes with documented activity relevant to cancer. Violin plots: the profiles show fitted density functions cropped at the minimal and maximal values; the internal horizontal lines show the upper quartile, median and lower quartile. DNA break data: gene DNA break counts were calculated within 5 kilobases downstream of the TSS (transcription start site) considering both strands; these counts were averaged across biological replicates.

## Notes

### Competing Interest Statement

The authors have declared no competing interest.

### Summary of Updates

Improved formatting - in pdf displayed on biorxiv, the biorxiv header overlapped with some of the figures and text

## REFERENCES

1. Rottenberg, S., Disler, C. & Perego, P. The rediscovery of platinum-based cancer therapy. Nature Reviews Cancer 21, 37–50 (2021).

2. da Costa, A.A.B.A., Chowdhury, D., Shapiro, G.I., D’Andrea, A.D. & Konstantinopoulos, P.A. Targeting replication stress in cancer therapy. Nature Reviews Drug Discovery 22, 38–58 (2023).

3. Larsen, A.K., Galmarini, C.M. & D’Incalci, M. Unique features of trabectedin mechanism of action. Cancer Chemother Pharmacol 77, 663–71 (2016).

4. Wang, J. et al. Trabectedin in Cancers: Mechanisms and Clinical Applications. Curr Pharm Des 28, 1949–1965 (2022).

5. Takebayashi, Y. et al. Antiproliferative activity of ecteinascidin 743 is dependent upon transcription-coupled nucleotide-excision repair. Nat Med 7, 961–6 (2001).

6. Olivieri, M. et al. A Genetic Map of the Response to DNA Damage in Human Cells. Cell 182, 481–496 e21 (2020).

7. Sakai, R., Rinehart, K.L., Guan, Y. & Wang, A.H. Additional antitumor ecteinascidins from a Caribbean tunicate: crystal structures and activities in vivo. Proc Natl Acad Sci U S A 89, 11456–60 (1992).

8. Pommier, Y. et al. DNA sequence-and structure-selective alkylation of guanine N2 in the DNA minor groove by ecteinascidin 743, a potent antitumor compound from the Caribbean tunicate Ecteinascidia turbinata. Biochemistry 35, 13303–9 (1996).

9. Moore, B.M., Seaman, F.C. & Hurley, L.H. NMR-Based Model of an Ecteinascidin 743−DNA Adduct. Journal of the American Chemical Society 119, 5475–5476 (1997).

10. Damia, G. et al. Unique pattern of ET-743 activity in different cellular systems with defined deficiencies in DNA-repair pathways. Int J Cancer 92, 583–8 (2001).

11. Marteijn, J.A., Lans, H., Vermeulen, W. & Hoeijmakers, J.H. Understanding nucleotide excision repair and its roles in cancer and ageing. Nat Rev Mol Cell Biol 15, 465–81 (2014).

12. Schärer, O.D. Nucleotide excision repair in eukaryotes. Cold Spring Harb Perspect Biol 5, a012609 (2013).

13. van den Heuvel, D., van der Weegen, Y., Boer, D.E.C., Ogi, T. & Luijsterburg, M.S. Transcription-Coupled DNA Repair: From Mechanism to Human Disorder. Trends Cell Biol 31, 359–371 (2021).

14. Zewail-Foote, M. & Hurley, L.H. Ecteinascidin 743: a minor groove alkylator that bends DNA toward the major groove. J Med Chem 42, 2493–7 (1999).

15. Seaman, F.C. & Hurley, L.H. Molecular Basis for the DNA Sequence Selectivity of Ecteinascidin 736 and 743: Evidence for the Dominant Role of Direct Readout via Hydrogen Bonding. Journal of the American Chemical Society 120, 13028–13041 (1998).

16. Zewail-Foote, M. & Hurley, L.H. Differential Rates of Reversibility of Ecteinascidin 743−DNA Covalent Adducts from Different Sequences Lead to Migration to Favored Bonding Sites. Journal of the American Chemical Society 123, 6485–6495 (2001).

17. Feuerhahn, S. et al. XPF-dependent DNA breaks and RNA polymerase II arrest induced by antitumor DNA interstrand crosslinking-mimetic alkaloids. Chem Biol 18, 988–99 (2011).

18. Jaspers, N.G. et al. Anti-tumour compounds illudin S and Irofulven induce DNA lesions ignored by global repair and exclusively processed by transcription-and replication-coupled repair pathways. DNA Repair (Amst) 1, 1027–38 (2002).

19. Tanasova, M. & Sturla, S.J. Chemistry and biology of acylfulvenes: sesquiterpene-derived antitumor agents. Chem Rev 112, 3578–610 (2012).

20. van Midwoud, P.M. & Sturla, S.J. Improved efficacy of acylfulvene in colon cancer cells when combined with a nuclear excision repair inhibitor. Chem Res Toxicol 26, 1674–82 (2013).

21. Otto, C. et al. Modulation of Cytotoxicity by Transcription-Coupled Nucleotide Excision Repair Is Independent of the Requirement for Bioactivation of Acylfulvene. Chem Res Toxicol 30, 769–776 (2017).

22. Ngo, L.P. et al. Sensitive CometChip assay for screening potentially carcinogenic DNA adducts by trapping DNA repair intermediates. Nucleic Acids Res 48, e13 (2020).

23. Staresincic, L. et al. Coordination of dual incision and repair synthesis in human nucleotide excision repair. EMBO J 28, 1111–20 (2009).

24. Hanasoge, S. & Ljungman, M. H2AX phosphorylation after UV irradiation is triggered by DNA repair intermediates and is mediated by the ATR kinase. Carcinogenesis 28, 2298–304 (2007).

25. Matsumoto, M. et al. Perturbed gap-filling synthesis in nucleotide excision repair causes histone H2AX phosphorylation in human quiescent cells. J Cell Sci 120, 1104–12 (2007).

26. Biggerstaff, M., Szymkowski, D.E. & Wood, R.D. Co-correction of the ERCC1, ERCC4 and xeroderma pigmentosum group F DNA repair defects in vitro. *EMBO J* **12**, 3685-92 (1993).

27. Wakasugi, M., Reardon, J.T. & Sancar, A. The non-catalytic function of XPG protein during dual incision in human nucleotide excision repair. J Biol Chem 272, 16030–4 (1997).

28. Sriramachandran, A.M. et al. Genome-wide Nucleotide-Resolution Mapping of DNA Replication Patterns, Single-Strand Breaks, and Lesions by GLOE-Seq. Mol Cell 78, 975–985 e7 (2020).

29. Seila, A.C. et al. Divergent Transcription from Active Promoters. Science 322, 1849–1851 (2008).

30. Preker, P. et al. RNA Exosome Depletion Reveals Transcription Upstream of Active Human Promoters. Science 322, 1851–1854 (2008).

31. Core, L.J., Waterfall, J.J. & Lis, J.T. Nascent RNA Sequencing Reveals Widespread Pausing and Divergent Initiation at Human Promoters. Science 322, 1845–1848 (2008).

32. Sigova, A.A. et al. Divergent transcription of long noncoding RNA/mRNA gene pairs in embryonic stem cells. Proceedings of the National Academy of Sciences 110, 2876–2881 (2013).

33. Wu, X. & Sharp, P.A. Divergent transcription: a driving force for new gene origination? Cell 155, 990–6 (2013).

34. Topka, S. et al. Targeting Germline-and Tumor-Associated Nucleotide Excision Repair Defects in Cancer. Clin Cancer Res 27, 1997–2010 (2021).

35. Borcsok, J. et al. Identification of a Synthetic Lethal Relationship between Nucleotide Excision Repair Deficiency and Irofulven Sensitivity in Urothelial Cancer. Clin Cancer Res 27, 2011–2022 (2021).

36. Fagbemi, A.F., Orelli, B. & Schärer, O.D. Regulation of endonuclease activity in human nucleotide excision repair. DNA Repair (Amst) 10, 722–9 (2011).

37. Matsunaga, T., Mu, D., Park, C.H., Reardon, J.T. & Sancar, A. Human DNA repair excision nuclease. Analysis of the roles of the subunits involved in dual incisions by using anti-XPG and anti-ERCC1 antibodies. *J Biol Chem* **270**, 20862-9 (1995).

38. Svoboda, D.L., Taylor, J.S., Hearst, J.E. & Sancar, A. DNA repair by eukaryotic nucleotide excision nuclease. Removal of thymine dimer and psoralen monoadduct by HeLa cell-free extract and of thymine dimer by Xenopus laevis oocytes. J Biol Chem 268, 1931–6 (1993).

39. Moggs, J.G., Yarema, K.J., Essigmann, J.M. & Wood, R.D. Analysis of incision sites produced by human cell extracts and purified proteins during nucleotide excision repair of a 1,3-intrastrand d(GpTpG)-cisplatin adduct. J Biol Chem 271, 7177–86 (1996).

40. Costanzo, F. et al. Promoters of ASCL1-and NEUROD1-dependent genes are specific targets of lurbinectedin in SCLC cells. EMBO Mol Med 14, e14841 (2022).

41. Gnügge, R., Reginato, G., Cejka, P. & Symington, L.S. Sequence and chromatin features guide DNA double-strand break resection initiation. Molecular Cell 83, 1237–1250.e15 (2023).

42. Kim, M. et al. Two interaction surfaces between XPA and RPA organize the preincision complex in nucleotide excision repair. Proceedings of the National Academy of Sciences 119, e2207408119 (2022).

43. van der Weegen, Y. et al. The cooperative action of CSB, CSA, and UVSSA target TFIIH to DNA damage-stalled RNA polymerase II. Nature Communications 11, 2104 (2020).

44. Sabatella, M. et al. Repair protein persistence at DNA lesions characterizes XPF defect with Cockayne syndrome features. Nucleic Acids Research 46, 9563–9577 (2018).

45. Ribeiro-Silva, C. et al. Ubiquitin and TFIIH-stimulated DDB2 dissociation drives DNA damage handover in nucleotide excision repair. Nature Communications 11, 4868 (2020).

46. Ge, J. et al. CometChip: a high-throughput 96-well platform for measuring DNA damage in microarrayed human cells. J Vis Exp, e50607 (2014).

47. Andrews, S. FastQC: a quality control tool for high throughput sequence data. (Babraham Bioinformatics, Babraham Institute, Cambridge, United Kingdom, 2010).

48. Bolger, A.M., Lohse, M. & Usadel, B. Trimmomatic: a flexible trimmer for Illumina sequence data. Bioinformatics 30, 2114–2120 (2014).

49. Langmead, B. & Salzberg, S.L. Fast gapped-read alignment with Bowtie 2. Nature Methods 9, 357–359 (2012).

50. Smith, T., Heger, A. & Sudbery, I. UMI-tools: modeling sequencing errors in Unique Molecular Identifiers to improve quantification accuracy. Genome Res 27, 491–499 (2017).

51. Li, H. et al. The Sequence Alignment/Map format and SAMtools. Bioinformatics 25, 2078–2079 (2009).

52. Quinlan, A.R. & Hall, I.M. BEDTools: a flexible suite of utilities for comparing genomic features. Bioinformatics 26, 841–842 (2010).

53. Ibarra, A., Benner, C., Tyagi, S., Cool, J. & Hetzer, M.W. Nucleoporin-mediated regulation of cell identity genes. Genes Dev 30, 2253–2258 (2016).

54. Muthalagu, N. et al. BIM is the primary mediator of MYC-induced apoptosis in multiple solid tissues. Cell Rep 8, 1347–53 (2014).

